# Semi-rational Engineering of Terminal Deoxynucleotidyl Transferase for High-efficiency Enzymatic *de Novo* DNA Synthesis

**DOI:** 10.64898/2025.12.29.696827

**Authors:** Chengjie Zhang, Zhicong He, Zhengquan Gao, Chunxiao Meng, Wei Peng, Lei Du, Shengying Li

## Abstract

The 21st century has witnessed rapid advancements in synthetic biology, with DNA synthesis emerging as a foundational technology. Conventional phosphoramidite-based methods face significant limitations, including short DNA elongation lengths (<300 nt), hazardous chemical waste, and low stepwise incorporation efficiency. Enzymatic DNA synthesis using terminal deoxynucleotidyl transferase (TdT) offers a promising alternative, enabling kilobase-scale assembly with greater efficiency and minimal environmental impact. Here, we identified *Bos taurus* TdT (BtTdT) through UniProt database mining as a catalytically active scaffold for natural and 3′-modified dNTPs. Comprehensive characterization of BtTdT’s enzymatic properties—including pH, temperature, metal ion dependence, and substrate specificity—revealed its optimal conditions. Truncation of the BRCT domain generated variants with enhanced activity compared to wild-type BtTdT. Guided by AlphaFold3-predicted structural models, we engineered a quintuple mutant (M5: Bt15AA^R336L/K338G/L397M/E456S/D395G^) optimized for 3′-ONH_2_-dNTP incorporation. M5 exhibited 30-fold activity enhancement relative to the triple mutant M3 (Bt15AA^R336L/K338G/L397M^) and achieved stepwise incorporation efficiency exceeding 98% in *de novo* synthesis of 10-nt ssDNA, demonstrating its potential for scalable enzymatic DNA synthesis. This work establishes a rational framework for TdT engineering through rational domain truncation and computational design, showing potential toward industrial-scale enzymatic DNA manufacturing.

## INTRODUCTION

The next generation of DNA sequencing technologies ^1–5^, diverse gene editing and assembly tools such as CRISPR/Cas9 ^6, 7^, Red/ET ^8^–^10^, Golden Gate ^11^, and Gibson Assembly ^12^, as well as other biotechnologies such as DNA origami ^13, 14^ and DNA data storage ^15–17^, all rely on efficient DNA synthesis, including the synthesis of short primers, long genes, and even entire chromosomes/genomes ^18–22^. At present, DNA synthesis is predominantly based on the sophisticated chemical solid-phase synthesis approach, namely, the phosphoramidite method ^21^ that was first introduced in 1981 ^23^. Despite continual refinements, this method remains limited to synthesizing DNA strands up to 200-300 nucleotides in length, thus requiring additional assembly techniques to construct longer sequences ^24, 25^. The solid-phase DNA synthesis also has other disadvantages, such as the use of toxic reagents and production of hazardous waste, posing significant environmental concerns ^20^. Furthermore, depurination during the synthetic process could break the DNA strands, which could severely affect the synthetic efficiency ^26–28^.

To address these limitations, enzymatic DNA synthesis has been emerging. This kind of DNA synthesis is primarily realized by terminal deoxynucleotidyl transferase (TdT), a template-independent DNA polymerase discovered in the 1960s that indiscriminately adds dNTP to the 3′ end of single-strand DNA (ssDNA), thereby continuously extending the DNA strand ^29^. Biologically, TdT was found to participate in the formation of diverse immunoglobulins and T cell receptors via V(D)J recombination (variable (V), diversity (D), and joining (J) gene segments) ^30–32^. In recent years, TdT has been adapted to controlled DNA synthesis, having achieved average stepwise coupling efficiency exceeding 97%, making it a promising candidate to synthesize longer DNA sequences ^33–35^. As TdT is the core of enzymatic DNA synthesis, great research efforts on optimizing TdT enzymes have been made on improving their catalytic efficiency and accuracy. The main strategies for TdT engineering include constructing combinatorial or saturation mutagenesis libraries for high-throughput screening^36–38^, and structure-guided semi-rational design enabled by computational simulations^39^. Although these approaches have significantly improved TdT’s catalytic efficiency and thermal stability, major bottlenecks remain that limit the industrial application of the best-in-class engineered TdTs. The key challenges include: (i) incomplete conversion across all substrates while maintaining high synthesis speed and fidelity, (ii) TdT’s thermal stability that is still inferior to commercially available DNA polymerases used in PCR—higher operating temperatures would help prevent DNA secondary structure formation and enable enzyme recyclability, thereby reducing synthesis costs, and (iii) markedly reduced catalytic activity on DNA substrates prone to forming secondary structures^40–42^.

In this study, we investigated the enzymatic properties of a select number of TdTs and identified *Bos taurus* TdT (BtTdT) as the most active candidate for incorporating 3′-ONH_2_ reversible terminator dNTPs under the optimized temperature, pH, and divalent metal ion concentration. Enzyme engineering efforts including *N*-terminal truncation, site-directed mutagenesis, and semi-rational design using AlphaFold 3 (AF3) led to an optimal BtTdT mutant M5, showing the highest reported activity for 3′-ONH_2_-dNTP-based DNA synthesis to date. Complete conversions of all 16 initiator DNA (iDNA) sequences with different terminal dinucleotides using all four 3′-ONH_2_-dNTP were achieved by M5. While testing iDNAs with variable terminal trinucleotide sequences, M5 is active against almost all combinations except for two combinations (GTC + 3′-ONH_2_-dGTP or 3′-ONH_2_-dCTP). A stepwise DNA synthesis of ten nucleotides was achieved with a high yield of target product, demonstrating its great potential for precise and scalable enzymatic DNA synthesis.

## RESULTS AND DISCUSSION

### Phylogenetic Tree Analysis and Screening of TdTs

Through bioinformatics-based homologous domain searches in the UniProt database, 226 TdTs were selected to build a phylogenetic tree. The sequence similarities between any two TdTs of the analyzed proteins are higher than 40%. These TdTs originate from different vertebrate taxa including *Cyclostomata*, *Pisces* (*Chondrichthyes* and *Osteichthyes*), *Amphibia*, *Reptilia*, *Aves* and *Mammalia*. Since TdTs of different origins show varied polymerization activities ^36^, it is essential to make parallel comparisons. Thus, we selected 11 representative TdTs having a variety of sequence identities with BtTdT that ranges from 40.9% to 98.8% for *in vitro* enzymatic activity assays (Figure 1, Tables S1-S2). Their coding sequences were optimized, inserted into pETDuet-1 plasmid, and transformed into *Escherichia coli* JM109(DE3) strains. After heterologous expression and protein purification using Ni-NTA His-tag purification agarose, nine His-tagged recombinant TdTs with high homogeneity were acquired (Figure S1). Notably, AmTdT from *Ambystoma mexicanum* (O57486) and EbTdT from *Eptatretus burgeri* (A0A8C4QBD0) failed to yield soluble target proteins.

**Fig. 1.**
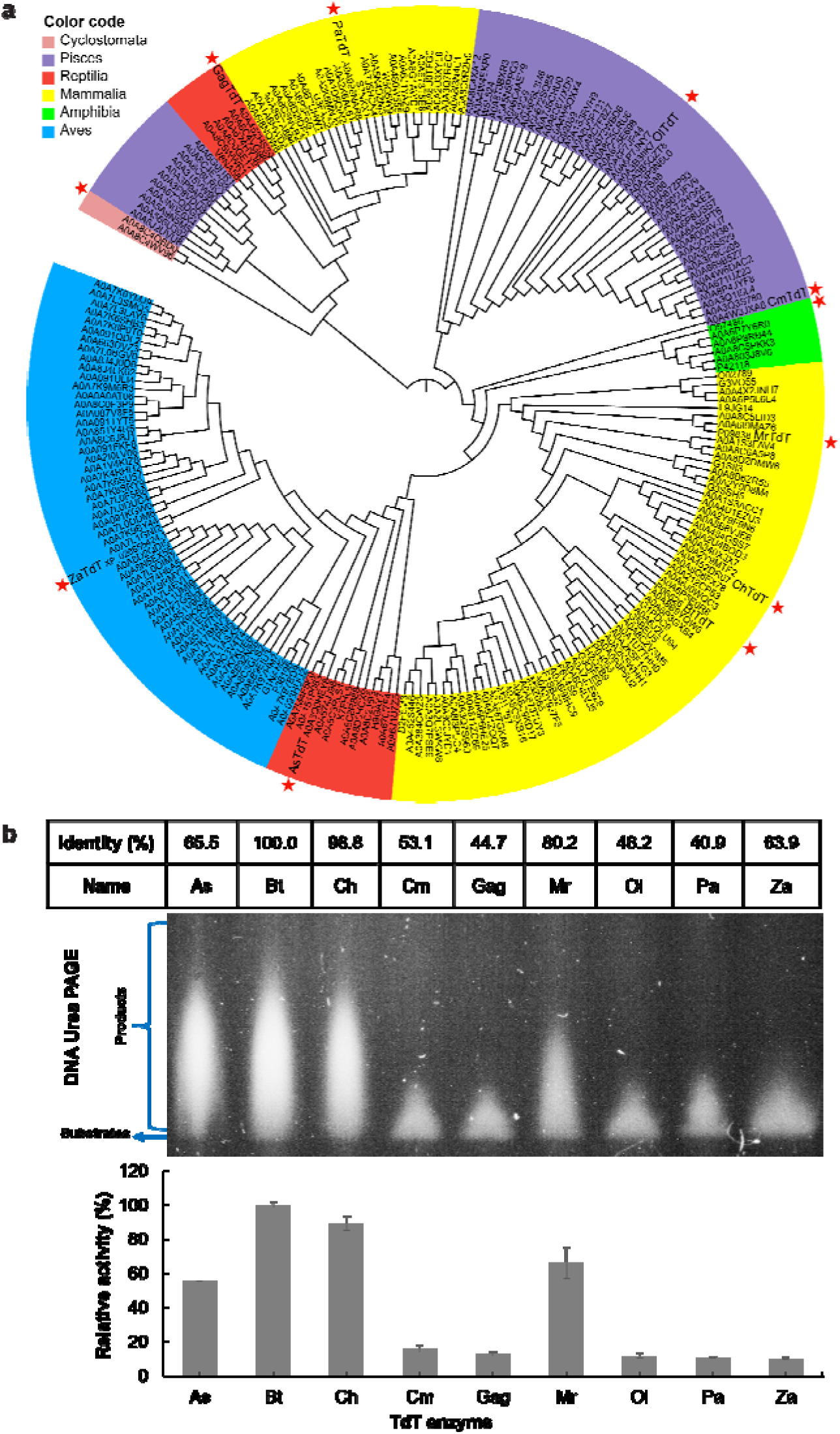
Evolutionary analysis and functional characterization of TdT enzymes. **a**, Phylogenetic analysis of 226 selected TdTs. **b**, Protein sequence identity (relative to BtTdT) and activity comparison (4 μM dT_20_ and 1 mM mixture of all four dNTPs (each at 0.25 mM) as substrates) of nine selected TdTs (as indicated by red stars in **a** and the following abbreviations). Species name abbreviation: *Ambystoma mexicanum* (Am), *Alligator sinensis* (As), *Bos taurus* (Bt), *Capra hircus* (Ch), *Callorhinchus milii* (Cm), *Eptatretus burgeri* (Eb), *Gopherus agassizii* (Ga), *Mus musculus* (Mm), *Oryzias latipes* (Ol), *Papio anubis* (Pa), and *Zonotrichia albicollis* (Za).

To make accurate activity comparisons, the gray values of each band on a DNA urea PAGE gel were quantified using ImageJ software^43^. Among the tested TdTs, BtTdT exhibited the highest activity towards natural dNTP substrates (Figure 1b). This result is inconsistent with the previous studies showing that ZaTdT exhibited higher activity than BtTdT^36, 39^. We attribute this inconsistency to the combined effects of extended reaction time, elevated assay temperature, and the use of all four native dNTPs, which collectively enabled a more comprehensive assessment of enzyme performance beyond initial velocity measurements. Further experiments showed that BtTdT also demonstrated significant catalytic activities towards the unnatural substrates 3′-ONH_2_-dNTP and 3′-OCN_3_-dNTP (Figure S2). The observation that both the P1-CC and P1-TC experimental groups displayed multiple distinct bands at different reaction times might result from TdT’s ability to remove a 3′-nucleotide from one oligonucleotide and transfer it to the 3′-end of another, a process known as the dismutase activity^44^. Based on these results, BtTdT was selected as the starting enzyme for subsequent studies on enzyme property characterization, protein engineering, and stepwise DNA synthesis.

### Catalytic Properties of BtTdT

To understand the enzyme properties of BtTdT, we evaluated its optimal temperature, pH, and metal ion cofactors. Specifically, the enzymatic activities of BtTdT gradually increased from pH 5.0 to 7.0, and then decreased from pH 7.0 to 9.0 (Figure 2a). Within the temperatures ranging from 20 to 50 °C, the highest activity was observed at 40 °C (Figure 2b), which was approximately 58% and 22% higher than that at 20 °C and 30 °C, respectively. To identity the most effective metal ion cofactor and corresponding concentration for BtTdT, 11 different types of monovalent and divalent metal ions were evaluated for their effects on enzyme activity. As a result, Co^2+^/Mg^2+^/Mn^2+^/Zn^2+^ showed positive effects on the polymerization activity (Figure 2c). Concentrations for these four metal ions were further investigated by testing the concentration range of 0-10 mM with a 2 mM interval. The optimal concentrations for Co^2+^, Mg^2+^, Mn^2+^, and Zn^2+^ were determined to be 2, 4, 4 and 2 mM, respectively (Figure 2d). Under combined optimal conditions at pH 7, 43 °C, and 1.5 mM Co^2+^, BtTdT exhibited a maximum activity (Figure 2e).

**Fig. 2.**
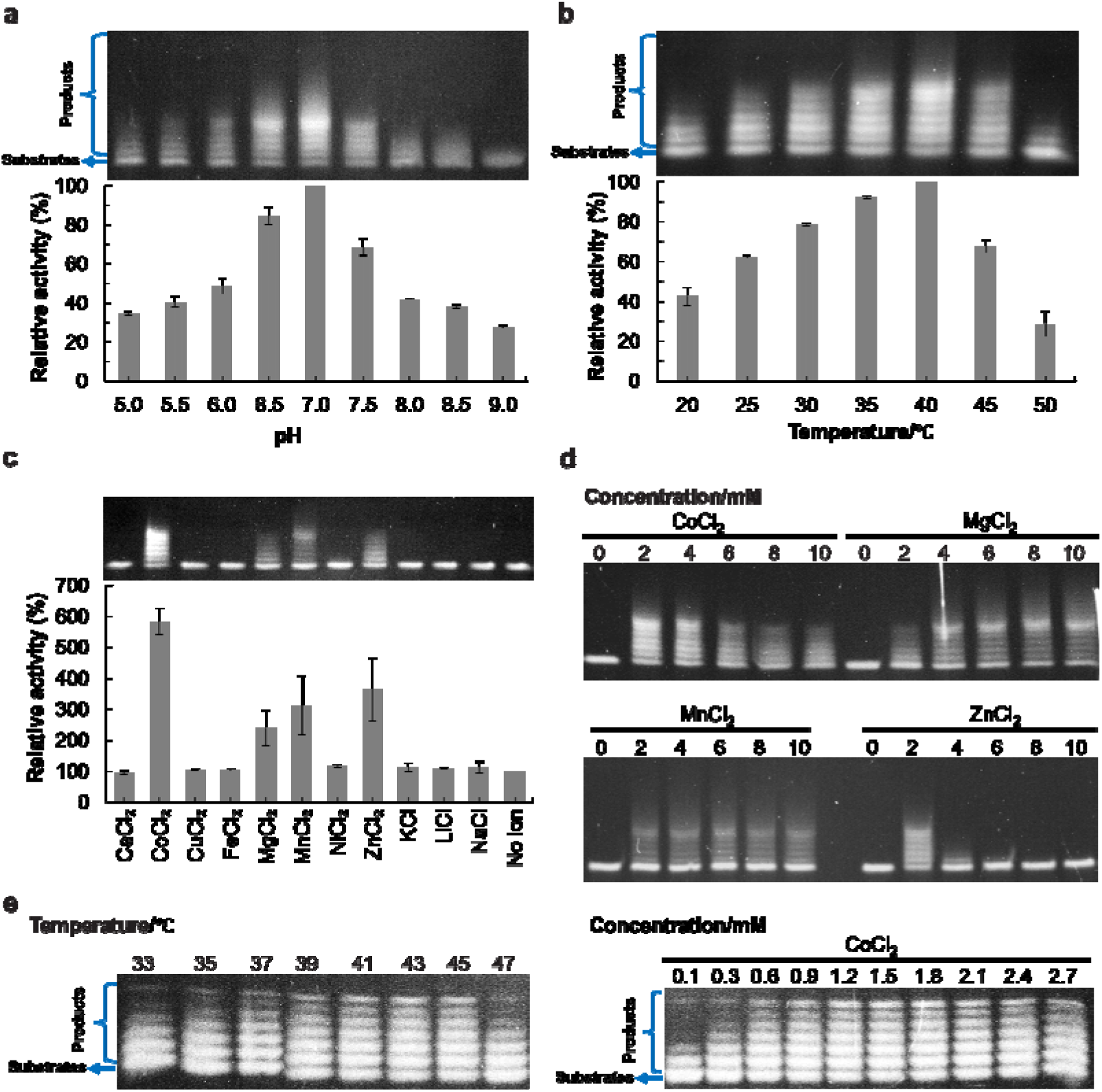
The activities of BtTdT under different reaction conditions. **a**, Optimal reaction pH of BtTdT. **b**, Optimal reaction temperature of BtTdT. **c**, Effects of different metal ions on BtTdT. **d**, Concentration gradients (0-10 mM) of four metal ions (Co^2+^/Mg^2+^/Mn^2+^/Zn^2+^) for BtTdT. **e**, More precise comparisons of the effects of temperature and Co^2+^ concentration on BtTdT activities.

### Truncation of BtTdT to Improve Catalytic Activity

The wild-type BtTdT comprises two structural domains: the BRCT domain and the catalytic domain POLXc. The POLXc domain consists of four subdomains, including the *N*-terminal α-helical subdomain (8 kDa domain), α-helical junction subdomain (finger domain), antiparallel β-sheet central subdomain (palm domain), and *C*-terminal subdomain (thumb domain) (Figure 3a). Previous studies indicated that removal of the BRCT domain while preserving the complete POLXc domain could enhance enzyme expression, stability, and DNA elongation activity ^39, 45^. Additionally, a conserved motif (residues 148-163) preceding the POLXc domain was found to be capable of stabilizing the catalytically active conformation through intramolecular interactions (*e.g.*, the interaction with Arg462 in the thumb domain) ^46^. To investigate the effect of the pre-POLXc motif on the enzymatic performance of BtTdT, we constructed a series of BRCT-deletion variants with incremental retention of the *N*-terminal residues prior to POLXc (Figure 3b). Enzymatic assays demonstrated that all truncated variants displayed a higher activity than the full-length BtTdT, with Bt15AA (residues 148-509, retaining 15 pre-POLXc amino acids, Figure 3b) showing the highest catalytic efficiency. This activity enhancement was consistently observed for both 3′-ONH_2_ and OCN_3_ modified dTTP substrates (Figure S3). Notably, variants with excessive truncations (Bt0AA, Bt2AA, and Bt5AA) showed poor solubility during protein preparation (Figure S4). Although the GST-tagged versions of Bt2AA and Bt5AA could be partially purified, both recombinant proteins exhibited low expression levels and lost their catalytic activity (Figure S4), corroborating the essential role of the 16-residue motif in maintaining enzyme functionality ^46^. These findings establish an effective truncation strategy for BtTdT engineering and provide mechanistic insights for future structure-guided optimization of TdTs.

**Fig. 3.**
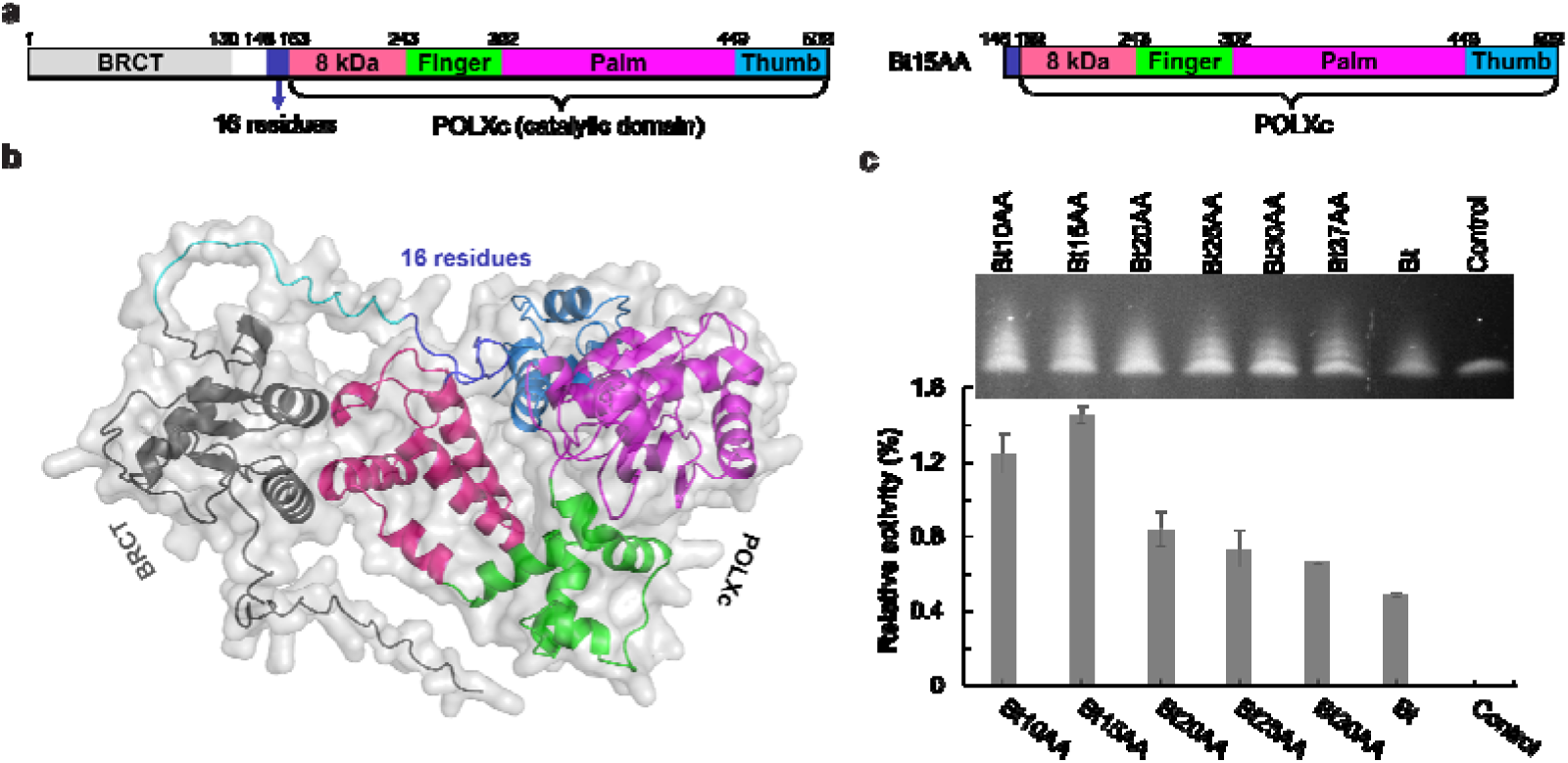
BtTdT and its truncation variants. **a**, Domain organization of intact BtTdT and the optimal truncation mutant Bt15AA. **b**, The three-dimensional structure of BtTdT (predicted by AF3) with all structural domains colored using the same color code in **a**. **c**, Catalytic activities of different BtTdT truncation mutants.

### Site-directed Mutagenesis of BtTdT

The catalytic performance of TdT could be influenced by three key factors: (i) the terminal dinucleotide composition of iDNA, (ii) natural dNTP substrate specificity, and (iii) compatibility with 3′-modified reversible for terminators (3′-ONH_2_, 3′-OCN_3_, and 3′-OC_2_CN dNTPs) ^33, 38, 39, 47^. To identify the optimal substrate for further engineering, we compared three classes of reversibly terminated dGTP analogues. Both wild-type BtTdT and the Bt15AA truncation mutant exhibited the highest activity toward 3′-ONH_2_-dGTP (Figure S2), thereby establishing this modified nucleotide as the preferred substrate for subsequent mutagenesis efforts.

To enhance catalytic efficiency, we performed structure-guided mutagenesis based on AF3-predicted three-dimensional models. Sequence alignment with MrTdT (*Mus musculus* TdT) revealed a conserved leucine residue (L397 in BtTdT, corresponding to L398 in MrTdT), the substitution of which with methionine was hypothesized to capable of improving base-stacking interactions between the 3′-end of iDNA and the incoming dNTP molecule ^36^. Our structural modeling confirmed the spatial proximity of L397 to the terminal nucleotide (Figure 4a and Figure S5), and the enzyme activity assays demonstrated that the Bt15AA^L397M^ mutant (M1) achieved a 1.5-fold increase in dNTP incorporation efficiency compared to Bt15AA (Figure 4b). For the unnatural substrate 3′-ONH_2_-dATP, M1 eliminated −1 nt byproducts compared with the wild-type enzyme (Figure S2), and its +1 nt activity was comparable to that of the wild-type enzyme.

**Fig. 4.**
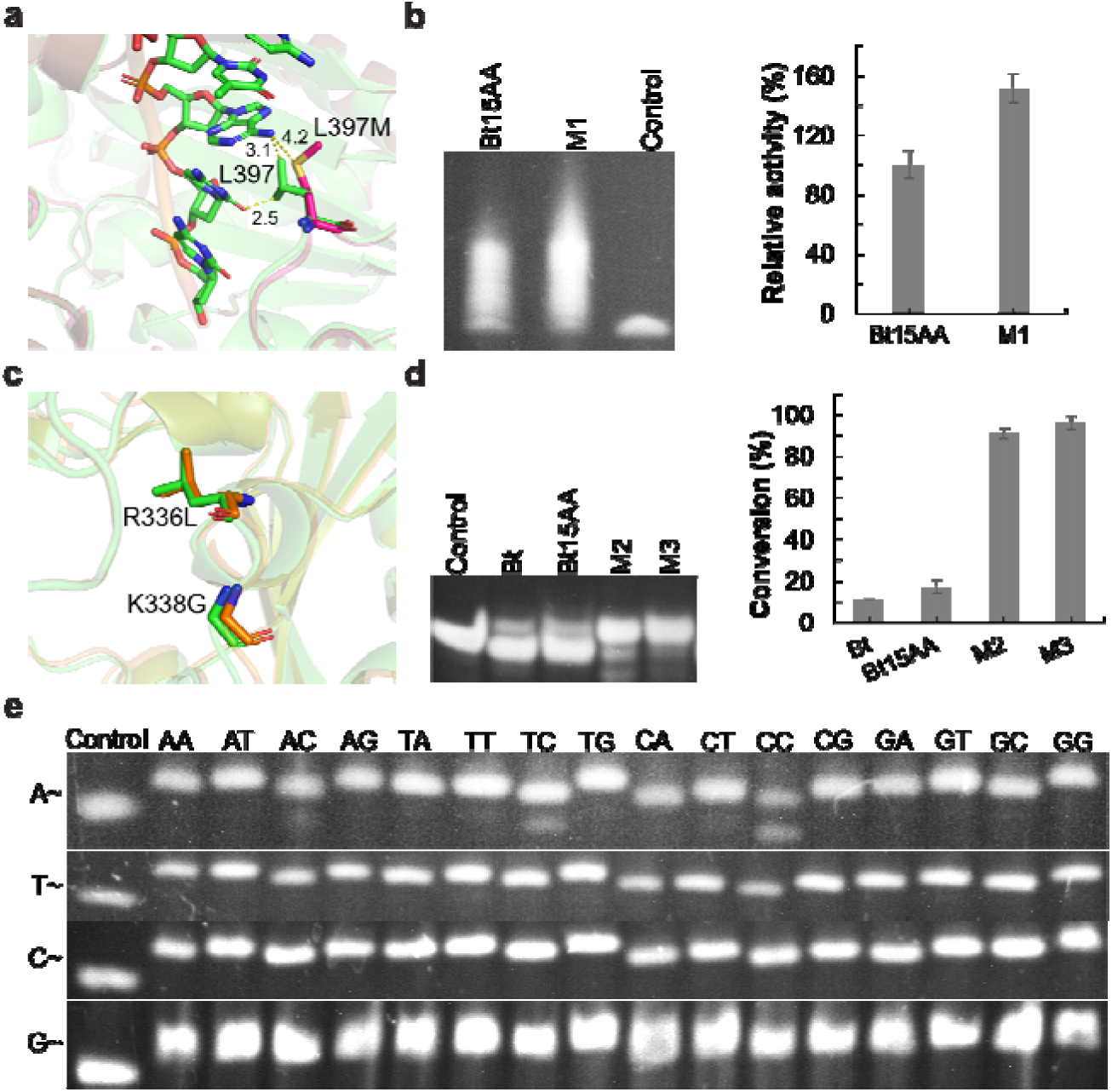
Mutagenesis analysis of BtTdT variants. **a**, L397 (green) or L397M (magenta) in BtTdT. **b**, Activity test for M1. **c**, R336/K338 in BtTdT (green) or R335/K337 in ZaTdT (orange). **d**, Activity comparison of three BtTdT mutants. **e**, Elongation by M3 with different iDNAs terminated with 16 dinucleotides and 4 types of 3′-ONH_2_-dNTPs. Reaction conditions: 1 mg/mL TdT, 1 µM iDNA, 42 °C, 10 min. “A∼, T∼, C∼, and G∼” stand for 3′-ONH_2_-dATP, 3′-ONH_2_-dTTP, 3′-ONH_2_-dCTP, and 3′-ONH_2_-dGTP, respectively. *Note*: Product bands at the same concentration may display different intensities due to sequence-specific effects.

Further structural insights from ZaTdT guided additional mutagenesis. The R336L/K338G double mutant of ZaTdT was previously reported to enhance 3′-ONH_2_-dNTP incorporation by enlarging the active site to better accommodate modified nucleotide conformations ^36^. Comparative structural analysis revealed conserved spatial positioning of residues R336 and K338 between BtTdT and ZaTdT (Figure 4c and Figure S5), suggesting that analogous substitutions in BtTdT might generate similar benefits. To our delight, this was confirmed experimentally that the Bt15AA^R336L/K338G^ mutant (M2) exhibited a 5-fold increase in activity than wild-type BtTdT (Figure 4d). Incorporation of the L397M mutation yielded the triple mutant Bt15AA^R336L/K338G/L397M^ (M3), which showed an additional 5% increase in catalytic efficiency with 3′-ONH_2_-dTTP compared to M2 (Figure 4d). Given the known influence of iDNA terminal dinucleotides on TdT activity ^47^, we further evaluated M3 against 16 iDNA substrates with varied terminal dinucleotide sequences and four types of 3′-ONH_2_-dNTPs. Remarkably, M3 achieved complete substrate conversion within 10 min for the majority of combinations. Exceptions were observed for TC- and CC-terminated iDNAs extended with 3′-ONH_2_-dATP, which only reached 70% and 50% conversions, respectively (Figure 4e).

The M3 variant demonstrated partial catalytic limitations, failing to achieve complete conversion of TC- or CC-terminated iDNAs with 3′-ONH_2_-dATP within 10 min (Figure 4e). In contrast, ZaTdT-R335L-K337G achieved full conversion of TC-terminated substrates under identical conditions (Figure S6), motivating the following structure-guided engineering to address this performance gap. Structurally, there are 21 amino acid residues in total within 5 Å of the catalytic center, which is defined as the bound dATP in the catalytic pocket. By aligning the three-dimensional structures of M5 and ZaTdT-R335L-K337G, we found that two out of these 21 aligned residues exhibit sequence divergence. Specifically, M3 contains Thr396 and Glu456 at the positions occupied by smaller residues (*i.e.*, Ala and Gly) in ZaTdT-R335L-K337G (Figure 5a). We reasoned that the bulkier side chains in M3 may create steric hindrance impeding 3′-ONH_2_-dATP incorporation. To mitigate this constraint, we designed four compensatory mutations including T396G, T396A, E456G, and E456A. As expected, all variants surpassed M3’s activity, with the E456G mutant achieving complete conversion of TC-terminated iDNA within 10 min and CC-terminated iDNA within 20 min (Figure 5b). However, combining Bt15AA^R336L/K338G/L397M/E456G^ with T396G or T396A (generating two quintuple mutants) conferred no additional benefit beyond that of the quadruple mutant Bt15AA^R336L/K338G/L397M/E456G^ (Figure 5b). Of note, Bt15AA^R336L/K338G/L397M/E456G^ maintained broad substrate compatibility, achieving complete conversions for 15 out of 16 iDNA types with 3′-ONH_2_-dATP (Figure 5c). And the persistent challenge of CC-terminated iDNA conversion (60% efficiency) still requires additional protein engineering efforts.

**Fig. 5.**
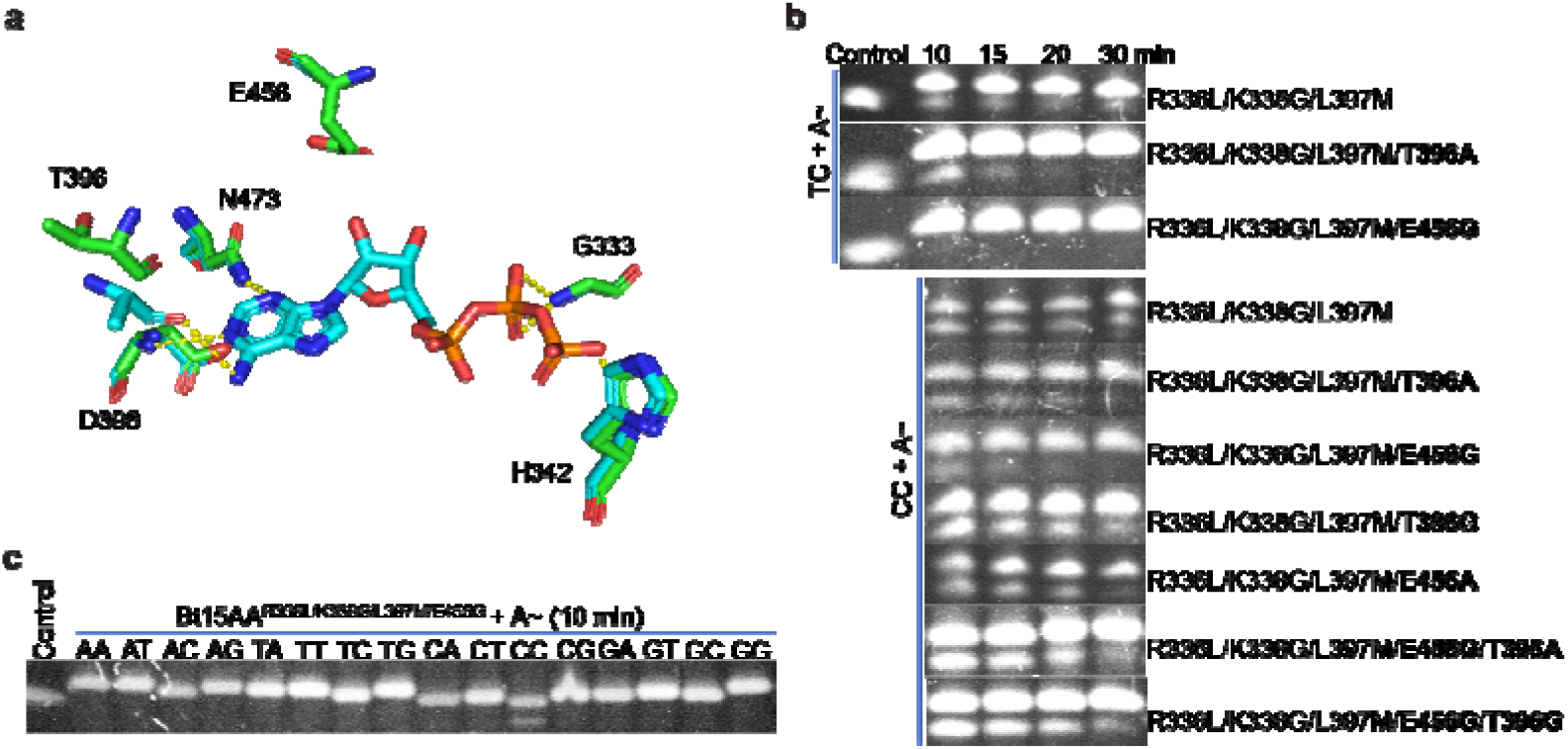
Further mutational work based on M3. **a**, Key amino acids at the catalytic center in BtTdT (green) and ZaTdT (cyan). **b**, Activity evaluation for M3-based mutants. **c**, Extension of 3′-ONH_2_-dATP by Bt15AA^R336L/K338G/L397M/E456G^. Reaction conditions: 1 mg/mL TdT, 1 µM iDNA, 42 °C, 10 min. *Note*: Product bands at the same concentration may display different intensities due to sequence-specific effects.

### Semi-rational Design Based on AlphaFold 3 Predicted Structures

Mechanistic insights into the critical role of residue E456 in 3′-blocked-dNTP incorporation efficiency prompted our further semi-rational engineering. We hypothesized that smaller amino acid substitution at this site would enlarge the catalytic pocket, thereby enhancing substrate flexibility and optimizing metal ion coordination geometry. To test this hypothesis, we employed AF3 for *in silico* modeling of twenty E456 variants of M3. Computational simulations incorporated ATP (a 3′-ONH_2_-dATP analog) and catalytic metal ions to mimic physiological binding states. Enzyme surface topologies were visualized using PyMOL. Because direct measurement of the enzyme surface-to-3′-end distance is impractical, changes in this distance in mutants relative to the wild-type—whether increased or decreased—were used to infer corresponding alterations in enzyme surface proximity. AF3 prediction revealed that E456 substitutions increase the spatial separation between the 3′-end of the substrate and enzyme surface, with E456A, E456G, and E456S mutants showing progressive increases of 0.3, 0.6, and 0.7 Å, respectively, when comparing to the wild-type enzyme (Figure 6a). Among these, E456S exhibits the most elevated position of the substrate sugar ring, suggesting a maximal reduction in steric hindrance and enhancement of conformational flexibility. We reasoned that the spatial reorganization would likely lead to a more stable and catalytically favorable substrate–metal interaction. Further experimental validation supported these computational predictions as the Bt15AA^R336L/K338G/L397M/E456S^ variant (M4) achieved complete conversion of all 16 iDNA types with 3′-ONH_2_-dATP within 5 min (Figure 6b). While testing the elongation time in 1 min with 3′-ONH_2_-dATP, the three iDNAs (terminated with TC, CT, or CC) could not be fully transformed. We further tested three other types of 3′-ONH_2_-dNTP substrates (16 iDNAs × 3 3′-ONH_2_-dNTPs). Results showed that M4 also demonstrated universal high efficiency across these substrates in 5-min reactions (Figure 6d). To gain mechanistic insight into enzyme specificity, iDNA P1-CC was substituted with P1-AC/TC/GC/AT/TT/CT/GT for structural simulations using AF3. The resulting models were then structurally aligned to identify deviations of the analog substrate ATP induced by iDNA in the catalytic pocket of M4 (Figure S7). For iDNA P1-NC, the 3′-OH of dATP corresponding to P1-TC/CC was positioned closest to the enzyme surface; and for iDNA P1-NT, the 3′-OH associated with P1-CT protruded nearest to the surface. These structural simulations were basically consistent with the observed activity profiles, suggesting that, in addition to key residue mutations that govern substrate dATP positioning in the catalytic pocket, the 3′-terminal base composition of the iDNA substrate may also influence substrate mononucleotide placement. However, the effect of the 3′-terminal base composition is relatively minor compared with that of key residue mutations. We speculate that this mechanistic insight may also apply to other engineered TdTs, such as ZaTdT or orthologs closely related to BtTdTs.

**Fig. 6.**
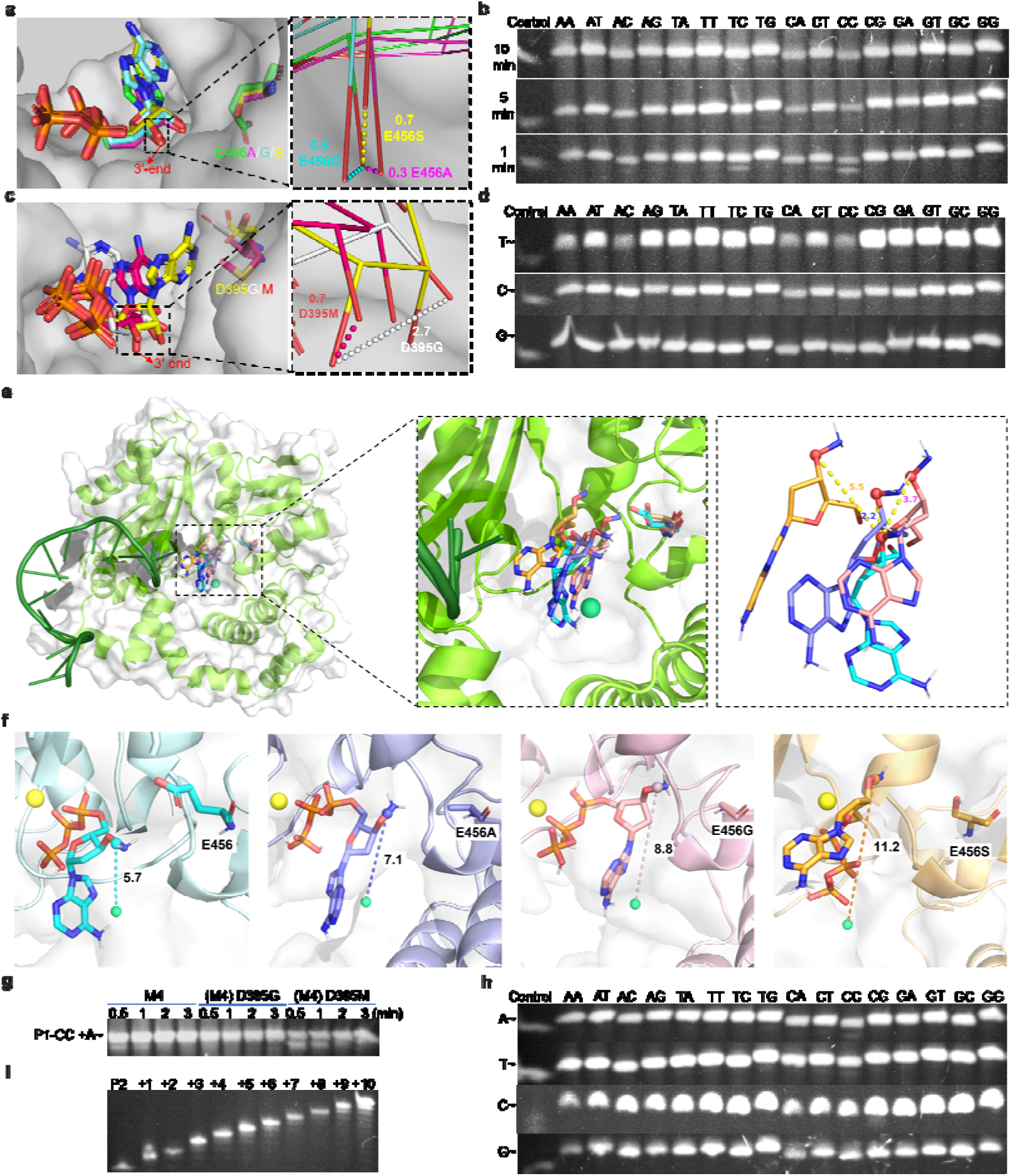
Mutagenesis based on AF3-based structural analysis. **a**, Position of surrogate substrate ATP in M3 variants with an amino acid substitution at E456. Enzyme-ligand complexes were predicted using AF3. E456S (yellow), E456G (aquamarine), E456 (green), and E456A (magenta). **b**, Activity test of M4 mutant after adding 3′-ONH_2_-dATP for different time. **c**, Position of surrogate substrate ATP in M4 variants with an amino acid substitution at D395. The distances in angstrom from the substrate’s 3′-end to the reference residues E456 or D395 (wild-type) are shown in different color for each mutant. Enzyme-ligand complexes were predicted using AF3. D395 (M4, yellow), D395G (white), and D395M (red). **d**, Extension of 3′-ONH_2_-dNTP by M4 in 5 min. Reaction conditions: 1 mg/mL TdT and 1 µM iDNA. **e**, Low-energy binding conformations of 3′-ONH_2_-dNTP in the WT and three E456 variants, along with their structural overlays. The distances between the 3′-O atom in the WT conformation and those in the mutant conformations are indicated. **f,** The average distance between the substrate’s O-NH_2_ moiety and the geometric center of the WT and three E456 mutant backbones. **g**, Time course assays of M4 mutations at D395. **h**, Extension of 3′-ONH_2_-dNTP by M5 in 30 s. Reaction conditions: 1 mg/mL BtTdT and 1 µM iDNA. **i**, PAGE analysis of the 10 steps of extension by M5. Extension conditions: 1 mg/mL M5, 1 µM iDNA P2, 1 min. “A∼, T∼, C∼, and G∼” stand for 3′-ONH_2_-dATP, 3′-ONH_2_-dTTP, 3′-ONH_2_-dCTP, and 3′-ONH_2_-dGTP, respectively. In panels **e** and **f,** blue, purple, pink, and orange denote wild type, E456A, E456G, and E456S, respectively. *Note*: Product bands at the same concentration may display different intensities due to sequence-specific effects.

To elucidate why the introduction of an NH_2_ group and the three E456 variants enhanced enzymatic activity, we performed molecular dynamics (MD) simulations using the substrate 3′-ONH_2_-dATP to examine its recognition in the wild-type (WT) and mutant enzymes. Our simulations revealed that, relative to the constructed starting conformation, 3′-ONH_2_-dATP readily undergoes conformational flipping, indicating that the NH_2_ substitution decreases the stability of substrate binding. In the low-energy conformer overlays, the O atom at the E456A variant exhibits a displacement of 2.2 Å relative to the WT 3′-O atom, whereas the displacements increase to 3.7 Å and 5.5 Å in the E456G and E456S mutants, respectively (Figure 6e). These positional shifts suggest that the altered low-energy conformations of the substrate reduce steric hindrance within the active site. This trend is consistent with the AF3 simulation results. Accordingly, we quantified the distances between the substrate’s O–NH_2_ group and the geometric center of the enzyme, the substrate centroid and the geometric center of the enzyme, and the substrate centroid and the geometric center of the active-site residues (G195, G196, H205, D206, D208, D258, T259, M260, D261, G311, W312, T313, G314, and N336), respectively (Figure 6f and Figure S8). The results consistently demonstrated a progressive increase in this distance across the WT, E456A, E456G, and E456S enzymes. These findings indicate that the NH_2_ substitution at the 3′-O position enhances substrate conformational flexibility and weakens substrate–protein interactions, while mutations at E456 further shift the substrate away from the catalytic center. Collectively, these structural rearrangements enlarge the active-site cavity, thereby facilitating improved enzymatic activity.

To further enhance the catalytic efficiency, we next targeted residue D395 adjacent to T396, based on previous reports implicating it in dNTP binding preferences ^48^. AF3 structural modeling reveals significant 3′-end displacement in D395 mutants. Specifically, D395M increases the substrate-enzyme surface distance by 0.7 Å when compared to the wild-type enzyme, while D395G achieves a remarkable 2.7 Å separation (Figure 6c). This spatial reorganization suggests enhanced substrate freedom, particularly in D395G where the 3′-ONH_2_ group adopts the most solvent-exposed conformation. Further functional assays confirmed these structural predictions. M4-derived variants (Bt15AAR336L/K338G/L397M/E456S/D395G and Bt15AAR336L/K338G/L397M/E456S/D395M) displayed significantly improved catalytic activity. Both Bt15AA^R336L/K338G/L397M/E456S/395G^ and Bt15AA^R336L/K338G/L397M/E456S/D395M^ achieved full conversion of 16 iDNAs with 3′-ONH_2_-dATP in 3 min, while Bt15AA^R336L/K338G/L397M/E456S/D395G^ (M5) achieved complete conversion within 1 min (Figure 6g). Furthermore, we synthesized and purified the previously characterized ZaTdT variants^36, 37, 39^ and conducted direct activity comparisons with M5 for P1-CC adding 3′-ONH_2_-dATP (Figure S9). Notably, M5 and M7-8/LG achieved complete substrate conversion within 10 min, whereas ZaTdT-R335L-A193T-G337H-H478G (which exhibits higher activity than ZaTdT-R335L-K337G) converted ∼50% under identical reaction conditions. The M7-8/LG mutant also showed optimal soluble expression in *E. coli*, yielding approximately 12 mg of purified enzyme per liter of LB culture, whereas M5 and ZaTdT-R335L-A193T-G337H-H478G exhibited lower yields of 3.5 and 2.5 mg/L, respectively. To evaluate its full substrate scope, M5 was examined across all 64 combinations of 16 iDNAs and four types of 3′-ONH_2_-dNTPs. The M5 variant demonstrated universal substrate compatibility, achieving almost complete extension in 30-sec reactions (Figure 6h).

Substrate bias at the last three nucleotides of DNA initiators has been reported for different TdTs^38, 39, 47^. We tested 64 (4×4×4) distinct iDNAs with all possible nucleotide combinations at the last three positions for incorporation of 3′-ONH_2_-dNTPs (Figure S10). As a result, M5 achieved near-complete conversion for most substrates but showed reduced activity toward CTC/CCC/CGA/CGT/CGC + 3′-ONH_2_-dATP combinations, and exhibited no detectable activity for GTC + 3′-ONH_2_-dC/GTP combinations. Enzymatic assays further revealed divergent activities for GTC + 3′-ONH_2_-dC/GTP: the wild-type and M1 maintained catalytic turnover, whereas M3 and M4 completely lost activity (Figure S10). Previously reported variants ZaTdT/LG and M7−8/LG also displayed substantially attenuated catalytic activity toward GTC + 3′-ONH_2_-dGTP or 3′-ONH_2_-dCTP, likely due to the collective influence of multiple residues, including R336L and K338G.

When catalyzing the P1-CT + 3′-ONH_2_-dC and dGTP reactions, M5 exhibited a higher coupling rate than with P1-CC + 3′-ONH_2_-dATP, achieving complete conversion within 2 and 3 min under identical conditions, respectively (Figure S11). Considering that the broad substrate promiscuity of an enzyme could enhance its application potential, we assessed M5’s activity toward other reversible nucleotide terminators, 3′-OCN_3_ and 3′-OC_2_CN dGTP. Activity profiling showed that M5 lost significant catalytic activity toward these substrates (Figure S12) relative to wild-type (Figure S2). The inability of M5 to broaden its substrate scope likely stems from the sterically extended side chains of these additional substrates (when compared with 3′-ONH_2_), rendering them incompatible with the NTP analog used in computational saturation mutagenesis.

Thermostability is often a significant challenge in TdT engineering, and trade-offs between enhanced activity and thermostability frequently occur during enzyme directed evolution. Thus, we measured the optimal reaction temperature of M5 (Figure S12). The temperature optimum of M5 (∼ 40 °C) coincided with that of wild-type BtTdT. Enzyme activity of M5 decreased sharply above 45 °C, indicating its limited thermostability. Further engineering will be required to enhance M5’s thermal tolerance to withstand process-favored temperatures, such as 60 °C, which can help minimize potential DNA secondary structures.

We also analyzed the universality of our mutations to other TdT orthologs and the comparative structural features of BtTdT in a wider phylogenetic context (Figure S13). R336 and K338 are strictly conserved across TdT orthologs, whereas the other three catalytic residue positions exhibit moderate sequence variation. Notably, the residues corresponding to R336 and K338 in previously reported hyperactive variants (ZaTdT-R335L-K337G, ZaTdT-R335L-A193T-G337H-H478G, and M7-8/LG)^36, 37, 39^ were also mutated, highlighting their functional indispensability for enhancing the catalytic activity toward 3′-ONH_2_-dNTP substrates.

### Mechanism Analysis of Improved Mutants

The remarkable activity enhancement of M5 relative to M3 (extension time reduced from > 30 min to 30 s) prompted mechanistic investigation through comparative structural analysis of the serial mutants including Bt15AA, M3, M4, and M5. To identify the structural determinants of the catalytic enhancement, we focused on the P1-CC/ATP substrate pair, a challenging combination that exhibited incomplete conversion by Bt15AA and M3 (> 30 min) and only moderate efficiency by M4 (Figure 5b and 6b). Comparative structural analysis revealed significant alterations in the interactions between the substrate and surrounding residues. Specifically, the number of substrate-interacting residues decreases from nine in Bt15AA to six in M5 (Figure 7a-7d), while the number of polar contacts such as hydrogen bonds and ionic interactions is reduced from nine to five. Similarly, the number of 5′-phosphate coordination bonds dramatically declines from seven to two. Importantly, the core phosphate-binding motifs (G341/H342 and G332/G333) remain conserved, thus preserving essential catalytic functions. This simplification of molecular interactions likely contributes to faster catalytic turnover through two synergistic mechanisms: (1) destabilizing pyrophosphate (PPi) retention post-cleavage to expedite product release ^49^, and (2) enabling 3′-ONH_2_-dNTP reorientation for optimal nucleophilic positioning and metal ion coordination (Mg^2+^/Mn^2+^ bridging), which is critical for transition-state stabilization. The inverse correlation between interaction complexity and enzymatic activity perhaps reflects an evolutionary trajectory of optimization through controlled flexibility-minimizing nonessential contacts to improve substrate dynamics without compromising the integrity of the catalytic core.

**Fig. 7.**
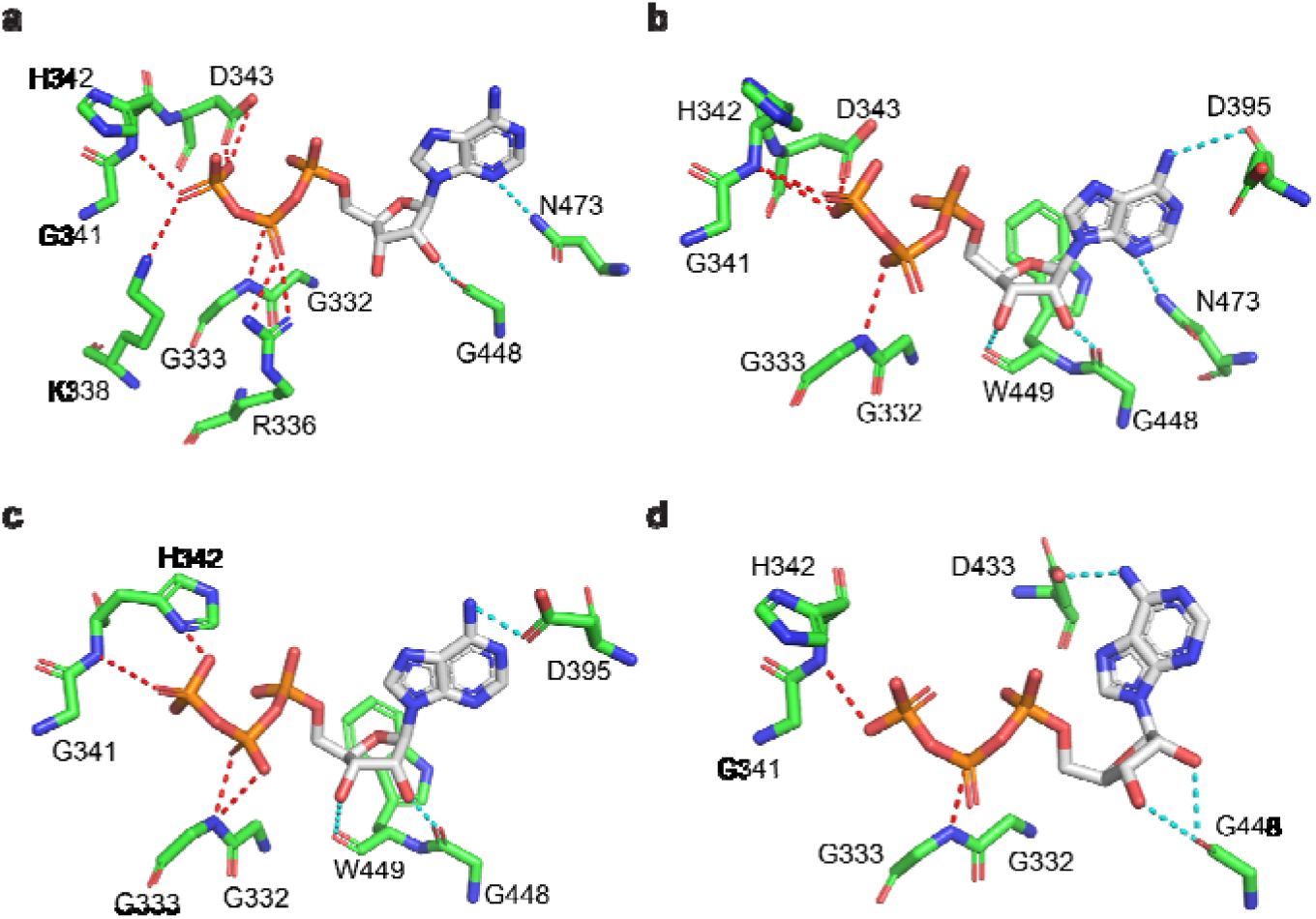
Mechanism analysis of improved mutants through enzyme-substrate interactions comparisons. **a**, The truncation mutant Bt15AA. **b**, M3 (Bt15AA^R336L/K338G/L397M^). **c**, M4 (Bt15AAR336L/K338G/L397M/E456S). **d**, M5 (Bt15AAR336L/K338G/L397M/E456S/D395G). The polar contacts between 5′-phosphate group of surrogate substrate ATP and surrounding amino acids are highlighted in red dashed lines. Amino acid residues are shown as green sticks, and substrate as white sticks.

### *De novo* DNA synthesis by Mutant M5

To evaluate the potential of M5 for *de novo* synthesis of long DNA sequences with 3′-ONH_2_-dNTPs as building blocks, a two-step oligonucleotide extension protocol was established to elongate an 18-nucleotide iDNA. We employed a streptavidin-biotin affinity system utilizing magnetic beads to capture the iDNA products, taking advantage of the high-affinity interactions between streptavidin and biotin-labeled iDNAs. The optimal BtTdT mutant M5 catalyzed DNA synthesis using 3′-ONH_2_-dNTPs as substrates in 1-min extension cycles. Each extension step was followed by a 1-min deprotection using 0.8 M sodium nitrite buffer (pH 5.2, adjusted using sodium acetate), which efficiently regenerated the natural 3′-OH terminus. By iteratively alternating between nucleotide incorporation and deprotection, full-length single-stranded DNA was synthesized *de novo*. After synthesis, DNA products were purified via magnetic separation and eluted for yield quantification. In a proof-of-concept demonstration, a 10-nucleotide ssDNA sequence (5′-ATGGATCCTC-3′) was synthesized achieving an overall cumulative yield of > 80% over 10 iterative cycles, with about 98% incorporation efficiency per step (0.8^1/10^) (Figure 6i).

## Conclusions

Although enzyme-mediated *de novo* DNA synthesis represents a transformative technology attracting global research interest, its practical application has been constrained by limited performance of TdT systems and inefficient synthetic workflows. In this work, through comparative analysis of natural TdTs, we identified BtTdT as a catalytically competent scaffold with significant engineering potential. Comprehensive characterization of BtTdT revealed critical limitations in substrate acceptance and reaction efficiency, motivating our structure-guided engineering efforts using 3′-ONH_2_-blocked nucleotides as substrates. Sequential introduction of mutations (Figure 8) effectively alleviated steric constraints on 3′-ONH_2_-dNTP binding, driving a progressive increase in substrate conversions and shifting reaction equilibria toward near-complete incorporation of 3′-ONH_2_-dNTPs. These results underscore that enhancing activity toward noncanonical substrates necessitates strategic disruption and reconfiguration of native substrate binding modes—rather than passive expansion of the binding pocket—to balance substrate mobility with precision alignment.

**Fig. 8.**
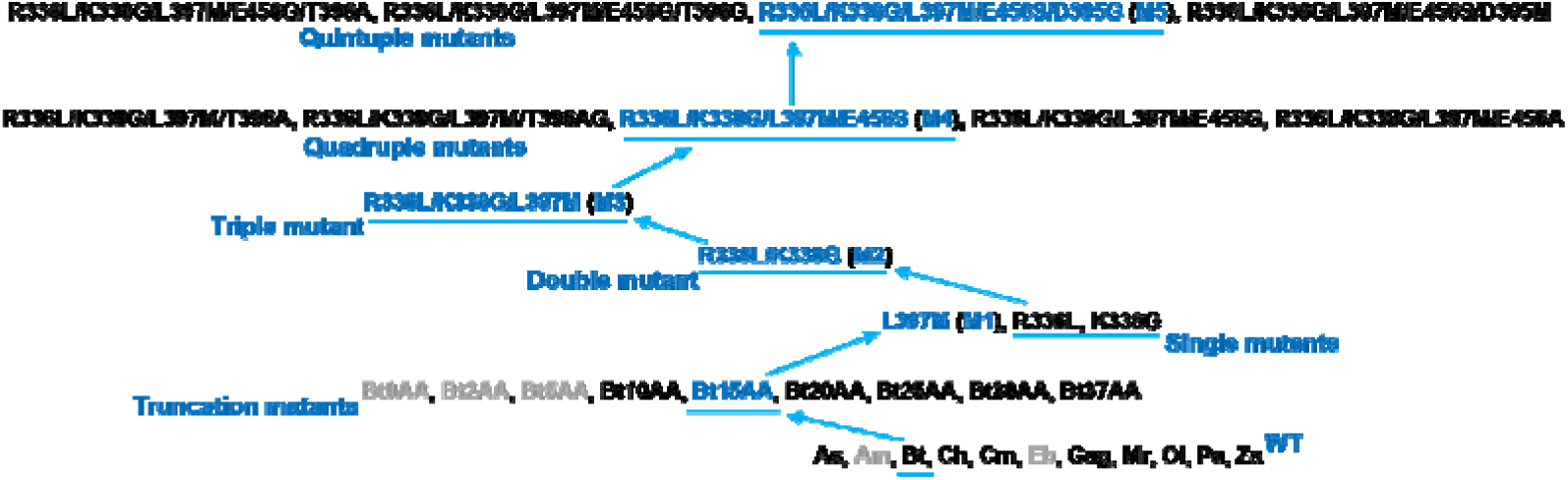
The summary of the TdT mutants generated in this study. The enzymes in grey stand for insoluble proteins.

The optimal mutant, Bt15AA^R336L/K338G/L397M/E456S/D395G^ (M5), demonstrated robust catalytic enhancement, enabling the *de novo* synthesis of medium-purity ssDNA of defined lengths (10 nt) with per-step incorporation efficiency exceeding 98%. Compared to its predecessor M3 (M3 and M5 exhibited conversion of ∼2.7% and 87% at 5 min, respectively), M5 exhibited a > 30-fold increase in catalytic efficiency with 3′-ONH_2_-dATP (Figure S11). This improvement was achieved through semi-rational design using AF3, which provided accurate structural predictions to guide substrate-binding site remodeling. The five key mutation sites identified in this work provide a rational blueprint for further engineering TdT activity toward both canonical and noncanonical dNTPs. This framework may also be applicable for other structurally homologous PolX family enzymes. For dNTPs with bulky 3’-blocking groups, combining saturation mutagenesis at these sites with MD simulations can accelerate the development of high-activity variants. This approach may particularly enable targeted enhancement of catalytic efficiency for specific and challenging iDNA/mononucleotide substrate combinations. Findings from our study and previous reports^33, 34, 36, 39^ lay the foundation for systematic screening and engineering TdT enzymes across broader orthologous families, with the aim of achieving high catalytic efficiency, broad substrate tolerance, and favorable expression profiles, thereby facilitating their potential application in industrial-scale DNA synthesis. This work highlights two key advances: a mechanistic model for TdT engineering through the controlled destabilization of nonproductive substrate interactions, and a predictive design framework that integrates AF3 to streamline the enzyme optimization process. These advances represent a significant step toward scalable, high-fidelity enzymatic DNA synthesis for industrial application.

## MATERIALS AND METHODS

### Strains, Media, and Chemicals

*Escherichia coli* DH5α (Beijing Tsingke Biotech, China) was used for plasmid construction and cloning. Protein expression was performed in either *E. coli* JM109 (DE3) or *E. coli* BL21 (DE3) strains (Shanghai Weidi Biotechnology, China). Luria-Bertani (LB) medium (0.5% yeast extract, 1% tryptone, and 1% NaCl) was used to culture *E. coli* strains for plasmids amplification and extraction, seed preparation, and protein expression and purification. The medium was supplemented with 50 mg/L ampicillin for plasmid maintenance, and protein expression was induced with 0.6 mM isopropyl β-D-1-thiogalactopyranoside (IPTG). SYBR^TM^ Gold nucleic acid gel stain was purchased from Thermo Fisher Scientific (USA) and 3’-ONH_2_-dNTPs were bought from Firebird Biomolecular Sciences (Florida, USA). Nitrous acid was acquired from Shanghai Yien Chemical Technology Co. Ltd. (Shanghai, China)Construction of Plasmids and Strains.

All TdT genes (see Table S5 in the Supporting Information) were codon-optimized for *E. coli* expression and synthesized by BGI Tech Solutions (Beijing Liuhe, Beijing, China). The strains, primers, and plasmids used or constructed in this study are shown in the Supporting Information. The TdT genes were cloned by PCR using PrimeSTAR Max DNA Polymerase (Takara Biomedical Technology, Beijing, China). Plasmids were constructed via homologous recombination using the ClonExpress Ultra One Step Cloning Kit purchased from Nanjing Vazyme Biotech (Nanjing, China). TdT genes were inserted into the pETDuet-1 plasmid to express proteins carrying the *N*-terminal His-tag. Vector pETDuet-1 was digested with restriction endonucleases *Bam*HI *and Not*I. The recombinant plasmids were transformed into cloning strains, and single colonies were picked and sent to Sangon Biotech (Qingdao, China) for sequencing. The recombinant plasmids were then transformed into *E. coli* JM109 (DE3) for protein expression and purification. Detailed procedures are provided in Table S3 and S4.

### Protein Expression and Purification of TdTs

*E. coli* JM109 (DE3) harboring certain recombinant vector was grown overnight in 6 mL LB medium with 100 µg/mL ampicillin at 37 °C with shaking at 220 rpm. The culture was used as a seed to inoculate 500 mL LB medium containing selective antibiotics and further grown at 37 °C, 220 rpm. When the culture optical density (OD_600_) reached approximately 0.6, the inducing temperature was adjusted to 18 °C, and IPTG was added to a final concentration of 0.6 mM to induce protein expression at 220 rpm for 22 h. Cells were harvested by centrifugation (5000 ×*g*, 10 min, 4 °C), and cell pellets were re-suspended in 60 mL precooled lysis buffer (50 mM Tris-HCl, 10% glycerol, pH 7.0). All purification steps were performed at 4 °C. First, an Ultrasonic Homogenizer (JY92-LDN) (Ningbo Scientz Biotechnology, China) was used for ultrasonication-based cell lysis (2 s on, 4 s off, and 30 min for total working time). After centrifugation at 10,000 ×*g* for 60 min to remove cell debris, the supernatant was transferred to a new 50 mL centrifuge tube. The recombinant His_6_-tagged proteins in the supernatant were purified using an Ni-NTA agarose column (Qiagen, Germany). The column was washed eight times with 10 mL lysis buffer with 50 mM imidazole, and the target proteins were eluted using a protein elution buffer (50 mM Tris-HCl, 10% glycerol, 250 mM imidazole, pH 7.0). An Amicon Ultra centrifugal filter with a 30 KDa cutoff (Merck KGaA, Germany) was used to concentrate the protein solution at 5,000 ×*g* for 30 min, and the elution buffer was exchanged with desalting buffer (100 mM Tris-HCl, 50% glycerol, pH 7.0) for three times. The concentration of the concentrated fraction (approximately 200 μL) was measured by absorbance spectrophotometry using a NanoDrop One^C^ spectrophotometer (Thermo Fisher Scientific, USA). Finally, protein aliquots (50 µL per tube) were flash-frozen in liquid nitrogen and stored at −80 °C for later use.

### Accurate Activity Measurement Based on Fluorescent Dye and Gray Value Calculations

**The** TdT enzymatic reactions for dNTPs were performed as follows: 0.3 μM purified TdTs, 4 μM iDNAs (P0; Table S4), 1 mM mixture of all four dNTPs (each at 0.25 mM), 2.5 mM CoCl_2_, and 100 mM Tris-HCl buffer (pH 7.0). Samples were mixed and incubated at 37 °C for 30 min to extend the iDNAs and then heated to 75 °C for 6 min to quench the reactions. The DNA products were analyzed on a 20% polyacrylamide gel with 8 M urea electrophoresis (DNA Urea PAGE), using a DNA Urea PAGE Electrophoresis Kit purchased from Real-Times Biotechnology (Beijing, China). The gel was stained with SYBR Gold nucleic acid gel dye and imaged on a Tanon 1600 Gel Image System (Tanon, China), and the products were quantitatively analyzed using ImageJ software based on gray value calculations.

### Determination of the Optimal Reaction Conditions for BtTdT (Temperature, pH, and Metal Ions)

The standard reaction system consisted of 0.3 μM purified BtTdT, 4 μM iDNAs (P0; Table S4), 1 mM dNTPs, 2.5 mM CoCl_2_, and 100 mM Tris-HCl buffer (pH 7.0). The samples were incubated at 37 °C for 30 min and at 75 °C for 6 min. For optimal temperature determination, the reaction system and stopping conditions were the same as mentioned above, and the enzyme reaction mix was aliquoted into different tubes. The tubes were then incubated at temperatures from 20 to 50 °C with an interval of 5 °C. For optimal pH determination, the reaction system and conditions were the same except for the pH level of 100 mM Tris-HCl buffer, which was set from 5.0 to 9.0, with a gap of 0.5. For the effects of metal ions on enzyme activity, the reaction system and conditions were the same apart from the kinds of metal ions used, and eleven kinds of metal ions were compared.

### Preparation of Substrate iDNAs for Magnetic-bead-based DNA synthesis

First, streptavidin magnetic beads (Dynabeads M-270 Streptavidin, Thermo Fisher Scientific Inc.) were prepared with wash buffer (10 mM Tris-HCl, 0.5 mM EDTA, 1 M NaCl, pH 7.5) to wash the beads twice and magnetic separation rack (Nanjing Vazyme Biotech, China) to separate the beads for 10 sec and then remove the supernatant. Beads were suspended with wash buffer (2 ×) and transferred into equivalent volume of biotin-labeled oligonucleotides sample dissolved by ultrapure water. The mixture was blended thoroughly and incubated using a rotary mixer HS-3 (Ningbo Scientz Biotechnology, China) at room temperature for 10-30 min or at 4 °C for 2 h. Then, streptavidin magnetic beads were separated from the mixtures by magnetic separation rack for 1 min, and washed twice with wash buffer. Biotin-labeled oligonucleotides were eluted with DNA/RNA elution buffer (95% formamide, 10 mM EDTA, pH8.2) at 65 °C for 5 min or 90 °C for 2 min.

iDNA strands elution condition was incubating samples in DNA/RNA elution buffer at 75-80 °C for 2 min, followed by magnetic separation for 30 sec and then transferring the supernatant to a new nuclease-free tube. The products were separated through magnetic rack separation method^50^ and washed twice to eliminate other impurities. The clean +1 products were put into next cycle to add another desired 3′-ONH_2_-dNTP with 1 min reaction time. Only one nucleotide was incorporated for each cycle, and after 10 cycles, the desired nucleotides were synthesized in a controlled manner.

### *De novo* DNA Synthesis of 10 nt ssDNA by Mutant M5

The 5′-biotinylated oligonucleotide P2 (Table S4) was immobilized on streptavidin magnetic beads as iDNA for *de novo* DNA synthesis. All nucleotide additions were performed at 42 °C for 1 min in EDS buffer, which contained 15% (*v*/*v*) DMSO, 50 mM *O*-benzylhydroxylamine HCl, 10% (*v*/*v*) glycerol, 0.05% (*v*/*v*) Tween 20, 2.5 mM tris-HCl, 40 mM NaCl, 0.5 mM Hepes, and 0.5 M cacodylic acid, with a final pH of 7.0 (measured in the absence of DMSO). Blocking groups of products 3′-ONH_2_-dNTPs were readily to deblock by treatment with 0.8 M sodium nitrite deblocking buffer at pH 5.2 (adjusted by sodium acetate) for 1 min and generated natural 3’-OH. For the 10 cycles of DNA (5′-ATGGATCCTC-3′) synthesis, the modified nucleotides were added in the following order for each synthetic step: (1) 3′-ONH_2_-dATP, (2) 3′-ONH_2_-dTTP, (3) 3′-ONH_2_-dGTP, (4) 3′-ONH_2_-dGTP, (5) 3′-ONH_2_-dATP, (6) 3′-ONH_2_-dTTP, (7) 3′-ONH_2_-dCTP, (8) 3′-ONH_2_-dCTP, (9) 3′-ONH_2_-dTTP, and (10) 3′-ONH_2_-dCTP.

### Enzyme-substrate interaction comparisons

Structural analysis of AlphaFold3-predicted mutant enzymes was performed using PyMOL to visualize substrate-binding interactions. The workflow included: (i) selecting the analog substrate ATP as the reference molecule, and (ii) identifying polar contacts between ATP and surrounding enzyme atoms. Enzyme–substrate interactions were subsequently visualized as dashed lines.

## Supporting information

The data that support the findings of this study are available in the supplementary material of this article.

## ASSOCIATED CONTENT

### Supporting Information

The Supporting Information is available free of charge at http://

## AUTHOR INFORMATION

### Author Contributions

S.L., L.D., C.M. and C.Z. designed the study and analyzed the data. C.Z. performed the experiments. Z.H. and W.P. performed the MD simulations. C.Z., C.M., L.D. and S.L. drafted the manuscript.

### Notes

The authors declare no competing financial interest.

## ACKNOWLEDGMENTS

This work was supported by National Key Research and Development Program of China (2021YFA0911500), Shandong Provincial Natural Science Foundation (ZR2020ZD23), National Natural Science Foundation of China (32370032, 32170088), the Taishan Young Scholars (tsqn202312032), the Fundamental Research Funds of Shandong University (2023QNTD001).

## REFERENCES

(1) Shendure, J.; Ji, H. Next-generation DNA sequencing. Nat. Biotechnol. 2008, 26 (10), 1135–1145.

(2) Metzker, M. L. Emerging technologies in DNA sequencing. Genome Res. 2005, 15 (12), 1767–1776.

(3) Hutter, D.; Kim, M. J.; Karalkar, N.; Leal, N. A.; Chen, F.; Guggenheim, E.; Visalakshi, V.; Olejnik, J.; Gordon, S.; Benner, S. A. Labeled nucleoside triphosphates with reversibly terminating aminoalkoxyl groups. Nucleos. Nucleot. Nucl. 2010, 29 (11), 879–895.

(4) Guo, J.; Xu, N.; Li, Z.; Zhang, S.; Wu, J.; Kim, D. H.; Sano Marma, M.; Meng, Q.; Cao, H.; Li, X.; Shi, S.; Yu, L.; Kalachikov, S.; Russo, J. J.; Turro, N. J.; Ju, J. Four-color DNA sequencing with 3’-*O*-modified nucleotide reversible terminators and chemically cleavable fluorescent dideoxynucleotides. Proc. Natl. Acad. Sci. U.S.A. 2008, 105 (27), 9145–9150.

(5) He, S.; Liu, Y.; Fang, S.; Li, Y.; Weng, T.; Tian, R.; Yin, Y.; Zhou, D.; Yin, B.; Wang, Y.; Liang, L.; Xie, W.; Wang, D. Solid-State nanopore DNA sequencing: Advances, challenges and prospects. Coord. Chem. Rev. 2024, 510, 215816.

(6) Sun, X.; Zhang, H.; Jia, Y.; Li, J.; Jia, M. CRISPR-Cas9-based genome-editing technologies in engineering bacteria for the production of plant-derived terpenoids. Eng. Microbiol. 2024, 4 (3), 100154.

(7) Cong, L.; Ran, F. A.; Cox, D.; Lin, S.; Barretto, R.; Habib, N.; Hsu, P. D.; Wu, X.; Jiang, W.; Marraffini, L. A.; Zhang, F. Multiplex genome engineering using CRISPR/Cas systems. Science 2013, 339 (6121), 819–823.

(8) Li, R.; Li, A.; Zhang, Y.; Fu, J. The emerging role of recombineering in microbiology. Eng. Microbiol. 2023, 3 (3), 100097.

(9) Muyrers, J. P.; Zhang, Y.; Stewart, A. F. Techniques: Recombinogenic engineering-new options for cloning and manipulating DNA. Trends Biochem. Sci. 2001, 26 (5), 325–331.

(10) Wang, H.; Li, Z.; Jia, R.; Hou, Y.; Yin, J.; Bian, X.; Li, A.; Müller, R.; Stewart, A. F.; Fu, J.; Zhang, Y. RecET direct cloning and Redαβ recombineering of biosynthetic gene clusters, large operons or single genes for heterologous expression. Nat. Protoc. 2016, 11 (7), 1175–1190.

(11) Engler, C.; Kandzia, R.; Marillonnet, S. A one pot, one step, precision cloning method with high throughput capability. PLoS One 2008, 3 (11), e3647.

(12) Gibson, D. G.; Young, L.; Chuang, R. Y.; Venter, J. C.; Hutchison, C. A, 3rd.; Smith, H. O. Enzymatic assembly of DNA molecules up to several hundred kilobases. Nat. Methods 2009, 6 (5), 343–345.

(13) Dey, S.; Fan, C.; Gothelf, K. V.; Li, J.; Lin, C.; Liu, L.; Liu, N.; Nijenhuis, M. A. D.; Saccà, B.; Simmel, F. C.; Yan, H.; Zhan, P. DNA origami. Nat. Rev. Methods Primers 2021, 1 (1), 13.

(14) Praetorius, F.; Kick, B.; Behler, K. L.; Honemann, M. N.; Weuster-Botz, D.; Dietz, H. Biotechnological mass production of DNA origami. Nature 2017, 552 (7683), 84–87.

(15) Church, G. M.; Gao, Y.; Kosuri, S. Next-generation digital information storage in DNA. Science 2012, 337 (6102), 1628.

(16) Yu, M.; Tang, X.; Li, Z.; Wang, W.; Wang, S.; Li, M.; Yu, Q.; Xie, S.; Zuo, X.; Chen, C. High-throughput DNA synthesis for data storage. Chem. Soc. Rev. 2024, 53 (9), 4463–4489.

(17) Weng, Z.; Li, J.; Wu, Y.; Xiu, X.; Wang, F.; Zuo, X.; Song, P.; Fan, C. Massively parallel homogeneous amplification of chip-scale DNA for DNA information storage (MPHAC-DIS). Nat. Commun. 2025, 16 (1), 667.

(18) Wu, Y.; Li, B. Z.; Zhao, M.; Mitchell, L. A.; Xie, Z. X.; Lin, Q. H.; Wang, X.; Xiao, W. H.; Wang, Y.; Zhou, X.; Liu, H.; Li, X.; Ding, M. Z.; Liu, D.; Zhang, L.; Liu, B. L.; Wu, X. L.; Li, F. F.; Dong, X. T.; Jia, B.; Zhang, W. Z.; Jiang, G. Z.; Liu, Y.; Bai, X.; Song, T. Q.; Chen, Y.; Zhou, S. J.; Zhu, R. Y.; Gao, F.; Kuang, Z.; Wang, X.; Shen, M.; Yang, K.; Stracquadanio, G.; Richardson, S. M.; Lin, Y.; Wang, L.; Walker, R.; Luo, Y.; Ma, P. S.; Yang, H.; Cai, Y.; Dai, J.; Bader, J. S.; Boeke, J. D.; Yuan, Y. J. Bug mapping and fitness testing of chemically synthesized chromosome X. Science 2017, 355 (6329), eaaf4706.

(19) Richardson, S. M.; Mitchell, L. A.; Stracquadanio, G.; Yang, K.; Dymond, J. S.; DiCarlo, J. E.; Lee, D.; Huang, C. L. V.; Chandrasegaran, S.; Cai, Y.; Boeke, J. D.; Bader, J. S. Design of a synthetic yeast genome. Science 2017, 355 (6329), 1040–1044.

(20) Hoose, A.; Vellacott, R.; Storch, M.; Freemont, P. S.; Ryadnov, M. G. DNA synthesis technologies to close the gene writing gap. Nat. Rev. Chem. 2023, 7 (3), 144–161.

(21) Ma, Y.; Zhang, Z.; Jia, B.; Yuan, Y. Automated high-throughput DNA synthesis and assembly. Heliyon 2024, 10 (6), e26967.

(22) Yu, W.; Dai, J.; Ma, Y. Recent development on DNA & genome synthesis. Curr. Opin. Syst. Biol. 2024, 37, 100490.

(23) Beaucage, S. L.; Caruthers, M. H. Deoxynucleoside phosphoramidites—A new class of key intermediates for deoxypolynucleotide synthesis. Tetrahedron Lett. 1981, 22 (20), 1859–1862.

(24) Caruthers, M. H. The chemical synthesis of DNA/RNA: our gift to science. J. Biol. Chem. 2013, 288 (2), 1420–1427.

(25) Li, H.; Huang, Y.; Wei, Z.; Wang, W.; Yang, Z.; Liang, Z.; Li, Z. An oligonucleotide synthesizer based on a microreactor chip and an inkjet printer. Sci. Rep. 2019, 9 (1), 5058.

(26) An, R.; Jia, Y.; Wan, B.; Zhang, Y.; Dong, P.; Li, J.; Liang, X. Non-enzymatic depurination of nucleic acids: factors and mechanisms. PLoS One 2014, 9 (12), e115950.

(27) LeProust, E. M.; Peck, B. J.; Spirin, K.; McCuen, H. B.; Moore, B.; Namsaraev, E.; Caruthers, M. H. Synthesis of high-quality libraries of long (150mer) oligonucleotides by a novel depurination controlled process. Nucleic Acids Res. 2010, 38 (8), 2522–2540.

(28) Deng, X.; Li, H.; Song, Y. Inkjet printing-based high-throughput DNA synthesis. Giant 2024, 17, 100222.

(29) Bollum, F. J. Calf thymus polymerase. J. Biol. Chem. 1960, 235 (8), 2399–2403.

(30) Desiderio, S. V.; Yancopoulos, G. D.; Paskind, M.; Thomas, E.; Boss, M. A.; Landau, N.; Alt, F. W.; Baltimore, D. Insertion of N regions into heavy-chain genes is correlated with expression of terminal deoxytransferase in B cells. Nature 1984, 311 (5988), 752–755.

(31) Jung, D.; Alt, F. W. Unraveling V(D)J recombination; insights into gene regulation. Cell 2004, 116 (2), 299–311.

(32) Boubakour-Azzouz, I.; Bertrand, P.; Claes, A.; Lopez, B. S.; Rougeon, F. Terminal deoxynucleotidyl transferase requires KU80 and XRCC4 to promote N-addition at non-V(D)J chromosomal breaks in non-lymphoid cells. Nucleic Acids Res. 2012, 40 (17), 8381–8391.

(33) Palluk, S.; Arlow, D. H.; de Rond, T.; Barthel, S.; Kang, J. S.; Bector, R.; Baghdassarian, H. M.; Truong, A. N.; Kim, P. W.; Singh, A. K.; Hillson, N. J.; Keasling, J. D. *De novo* DNA synthesis using polymerase-nucleotide conjugates. Nat. Biotechnol. 2018, 36 (7), 645–650.

(34) Verardo, D.; Adelizzi, B.; Rodriguez-Pinzon, D. A.; Moghaddam, N.; Thomée, E.; Loman, T.; Godron, X.; Horgan, A. Multiplex enzymatic synthesis of DNA with single-base resolution. Sci. Adv. 2023, 9 (27), eadi0263.

(35) Zhang, C.; Subthain, H.; Guo, F.; Fang, P.; Zheng, S.; Shen, M.; Jiang, X.; Gao, Z.; Meng, C.; Li, S.; Du, L. Terminal deoxynucleotidyl transferase: Properties and applications. Eng. Microbiol. 2025, 5 (1), 100179.

(36) Lu, X.; Li, J.; Li, C.; Lou, Q.; Peng, K.; Cai, B.; Liu, Y.; Yao, Y.; Lu, L.; Tian, Z.; Ma, H.; Wang, W.; Cheng, J.; Guo, X.; Jiang, H.; Ma, Y. Enzymatic DNA synthesis by engineering terminal deoxynucleotidyl transferase. ACS Catal. 2022, 12 (5), 2988–2997.

(37) Hu, L.; Zhang, Z.; Li, C.; Han, M.; Hao, M.; Zhang, X.; Ahmed, N.; Luo, J.; Lu, X.; Sun, J.; Jiang, H. High-Throughput Screening for Oligonucleotide Detection by ADE-OPI-MS. Analytical Chemistry 2024, 96 (29), 12040–12048.

(38) Forget, S. M.; Krawczyk, M. J.; Knight, A. M.; Ching, C.; Copeland, R. A.; Mahmoodi, N.; Mayo, M. A.; Nguyen, J.; Tan, A.; Miller, M.; Vroom, J.; Lutz, S. Evolving a terminal deoxynucleotidyl transferase for commercial enzymatic DNA synthesis. Nucleic Acids Res. 2025, 53 (4), gkaf115.

(39) Niu, Y.; Chen, B.; Zhang, H.; Zheng, W.; Wu, J.; Yang, L.; Yang, M.; Yu, H. Computational design of a thermostable and highly active terminal deoxynucleotidyl transferase for synthesis of long *de novo* DNA molecules. ACS Catal. 2025, 15 (9), 7201–7216.

(40) Fessner, N. D. Enzymatic Strategies for Next-Generation DNA Synthesis: Boosting Efficiency and Overcoming Secondary Structures. ACS Catal. 2025, 16106–16114.

(41) Gao, N.; Yu, A.; Yang, W.; Zhang, X.; Shen, Y.; Fu, X. Enzymatic *de novo* oligonucleotide synthesis: Emerging techniques and advancements. Biotechnol. Adv. 2025, 82, 108604.

(42) Zhou, H.; Li, H.; Jia, Z.; Niu, S.; Song, Y. Reaction pathways and technologies of *in vitro* DNA synthesis. Cell Rep. Phys. Sci. 2025, 6 (8), 102777.

(43) Schneider, C. A.; Rasband, W. S.; Eliceiri, K. W. NIH Image to ImageJ: 25 years of image analysis. Nat. Methods 2012, 9 (7), 671–675.

(44) Anderson, R. S.; Bollum, F. J.; Beattie, K. L. Pyrophosphorolytic dismutation of oligodeoxy-nucleotides by terminal deoxynucleotidyltransferase. Nucleic Acids Res. 1999, 27 (15), 3190–3196.

(45) Li, A.-N.; Shi, K.; Zeng, B.-B.; Xu, J.-H.; Yu, H.-L. Enhancing the expression of terminal deoxynucleotidyl transferases by N-terminal truncation. Biotechnol. J. 2024, 19 (9), 2400226.

(46) Delarue, M.; Boulé, J. B.; Lescar, J.; Expert-Bezançon, N.; Jourdan, N.; Sukumar, N.; Rougeon, F.; Papanicolaou, C. Crystal structures of a template-independent DNA polymerase: murine terminal deoxynucleotidyltransferase. EMBO J. 2002, 21 (3), 427–439.

(47) Schaudy, E.; Lietard, J.; Somoza, M. M. Sequence preference and initiator promiscuity for *de novo* DNA synthesis by terminal deoxynucleotidyl transferase. ACS Synth. Biol. 2021, 10 (7), 1750–1760.

(48) Ukladov, E. O.; Tyugashev, T. E.; Kuznetsov, N. A. Computational modeling study of the molecular basis of dNTP selectivity in human terminal deoxynucleotidyltransferase. Biomolecules 2024, 14 (8), 961.

(49) Gouge, J.; Rosario, S.; Romain, F.; Beguin, P.; Delarue, M. Structures of intermediates along the catalytic cycle of terminal deoxynucleotidyltransferase: dynamical aspects of the two-metal ion mechanism. J. Mol. Biol. 2013, 425 (22), 4334–4352.

(50) Haukanes, B.-I.; Kvam, C. Application of magnetic beads in bioassays. Nat. Biotechnol. 1993, 11 (1), 60–63.

