## Supplementary material for "Semi-rational Engineering of Terminal Deoxynucleotidyl Transferase for High-efficiency Enzymatic *de Novo* DNA Synthesis": The data that support the findings of this study are available in the supplementary material of this article.

The PDF file includes:

Figures S1 to S13

Tables S1 to S4

**Molecular Dynamic Simulation.**

In this study, MD simulations of the Co enzyme-DNA-substrate complexes were conducted utilizing the Amber Package, version 22 (University of California, SanFrancisco, CA)^1^ to investigate the effects of mutations at the 319 site (including E319A, E319G, and E319S) on the substrate binding modes within the active site. The initial conformations for the MD simulations of Co enzyme-DNA-substrate complexes were derived from the optimized structures obtained via *in silico* docking. The ff14SB^2^ and OL15^3^ force field were loaded in the “tleap” program to parameterize the amino acid residues within Co enzyme and DNA half-sites, respectively, while the GAFF force field^4^ was applied to parameterize the substrate. The simulation system was solvated with a periodic cubic box (the volume is 84.193×132.987×99.174 Å^3^) filled with TIP3P water molecules and an approximate number (13) of sodium counterion to neutralize the charge. In total, there are 28912 crystallographic solvent molecules, and the shortest distance between the complex and the box boundary is set to 10 Å.

After proper setup, the complex systems were successively fully minimized by combining 10,000 steps of the most rapid descent method and 10,000 steps of the conjugate gradient method, and each system was gradually heated from 0 K to 300 K for a total of 50 ps using the NVT ensemble. To achieve a uniform density after heating dynamics, 1 ns of density equilibrium was executed under the NPT ensemble, during which the temperature and pressure of the system were maintained at 300 K and 1.0 atm by utilizing the Langevin thermostat and the Berendsen barostat,^5,6^ respectively. Subsequently, all complexes were equilibrated for 2 ns without any restraints in order to relieve minor unfavorable ligand-protein steric interactions that might have been still present. Finally, a productive MD simulation of 100 ns was carried out under the NPT ensemble. Throughout the process, the simulation integration step was set to 2 fs, and the conformational configurations of complex systems were saved every 100 ps in trajectory files. The covalent bonds connecting hydrogen atoms were restricted with the SHAKE algorithm.^7^ The short-range nonbonded interactions were adopted using a cutoff radius of 10 Å, while the long-range electrostatic interactions were modeled utilizing the Particle mesh Ewald (PME) method with a grid point density of 0.1 nm and an interpolation order of 4.^8^ The results of MD simulations were presented by using the Visual Molecular Dynamics 1.9.3a,^9^ and the PyMOL software^10^ was used to exhibit the graphics of the MD simulations.

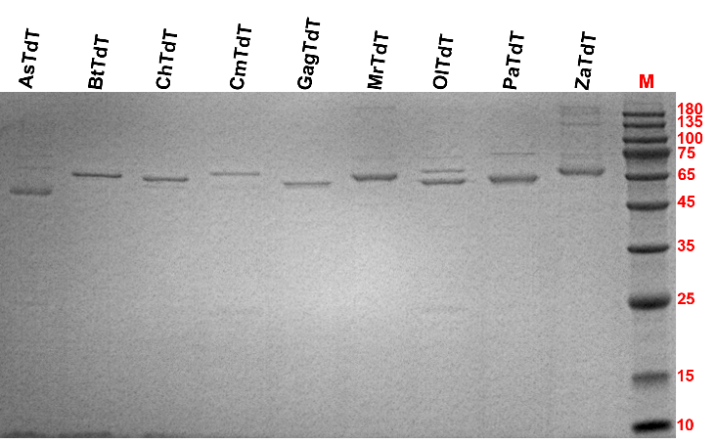

**Figure S1.** SDS-PAGE analysis of nine different purified TdTs.

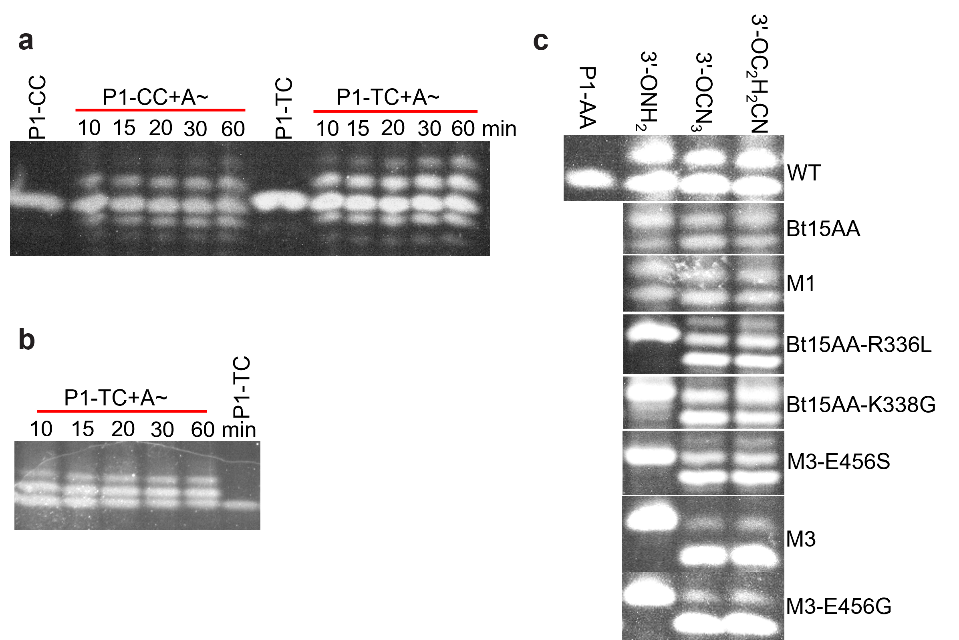

**Figure S2.** The incorporation of 3′-ONH_2_-dATP, 3′-ONH_2_/3′-OCN_3_/3′-OC_2_H_2_CN dGTPs for wild-type (**a**), M1 (**b**) and mutant BtTdTs (**c**). A~ stands for 3′-ONH_2_-dATP.

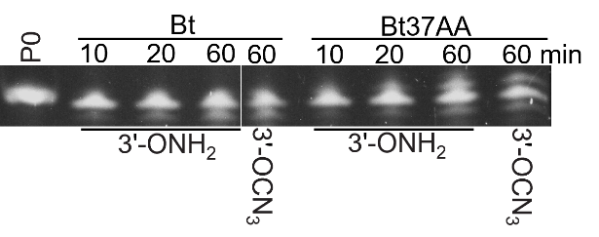

**Figure S3.** The activity test of Bt and Bt37AA for elongating 3′-ONH_2_ and OCN_3_ dTTP.

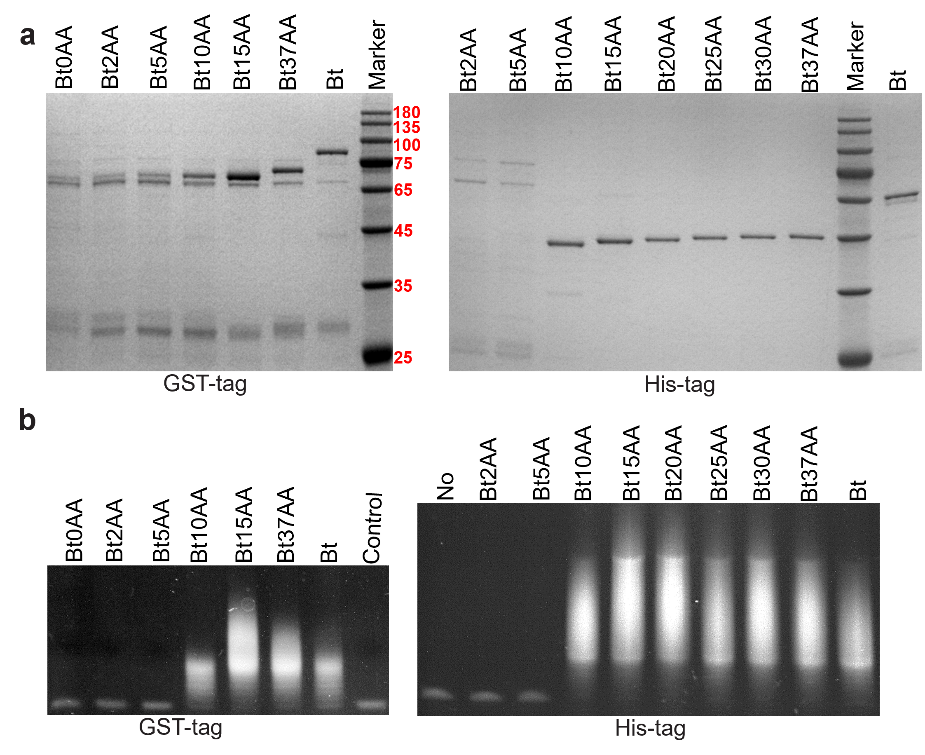

**Figure S4.** Purification of different truncation mutants of BtTdT with GST or His tag (**a**) and activity test of each mutant (**b**). The control group omitted enzyme.

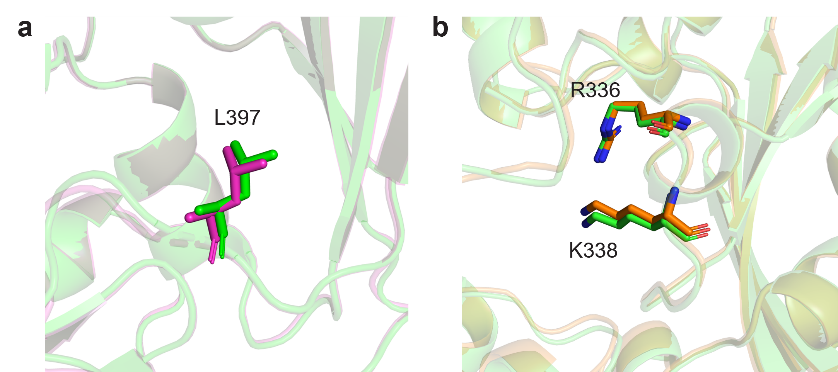

**Figure S5. a**, L397 in BtTdT (green) or L398 in MrTdT (magenta). **b**, R336/K338 in BtTdT (green) or R335/K337 in ZaTdT (orange).

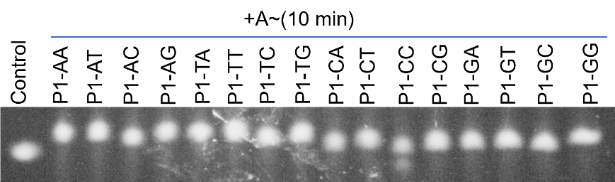

**Figure S6.** The elongation rate of ZaTdT-R335L-K337G for different initiators terminated with 16 kinds of dinucleotides and 3′-ONH_2_-dATP.

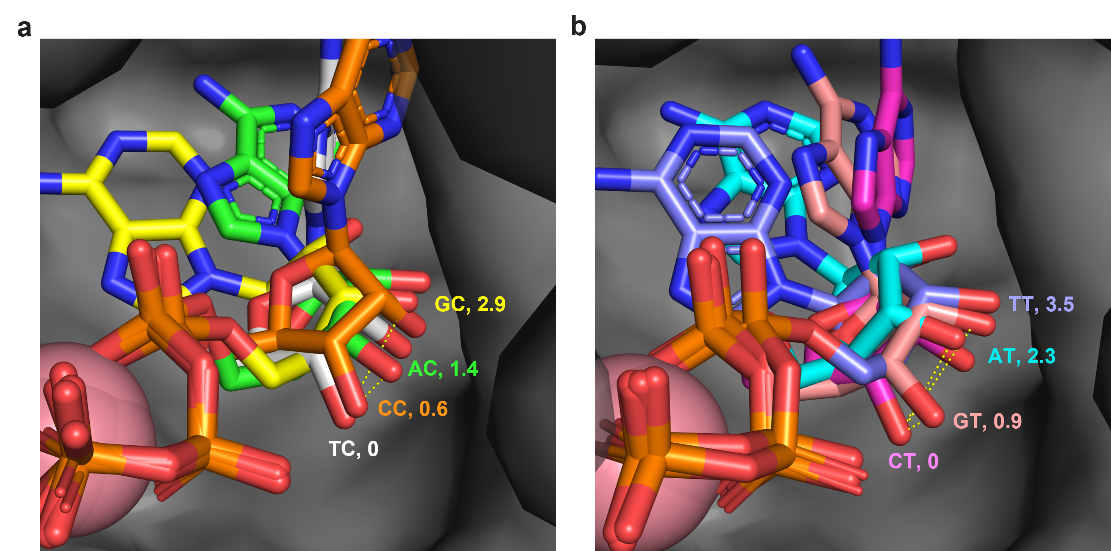

**Figure S7.** Position of surrogate substrate ATP in M4 with different iDNAs. Enzyme-ligand complexes were predicted using AF3. P1-TC (white), P1-CC (orange), P1-AC (green), P1-GC (yellow), P1-CT (light purple), P1-GT (pink), P1-AT (cyan), and P1-TT (light blue). The distances were numerically quantified between the 3'-OH of ATP in M4 for four distinct iDNAs and the 3'OH of the dATP closest to the enzyme surface (in **a**, P1-TC’; in **b**, P1-CT’).

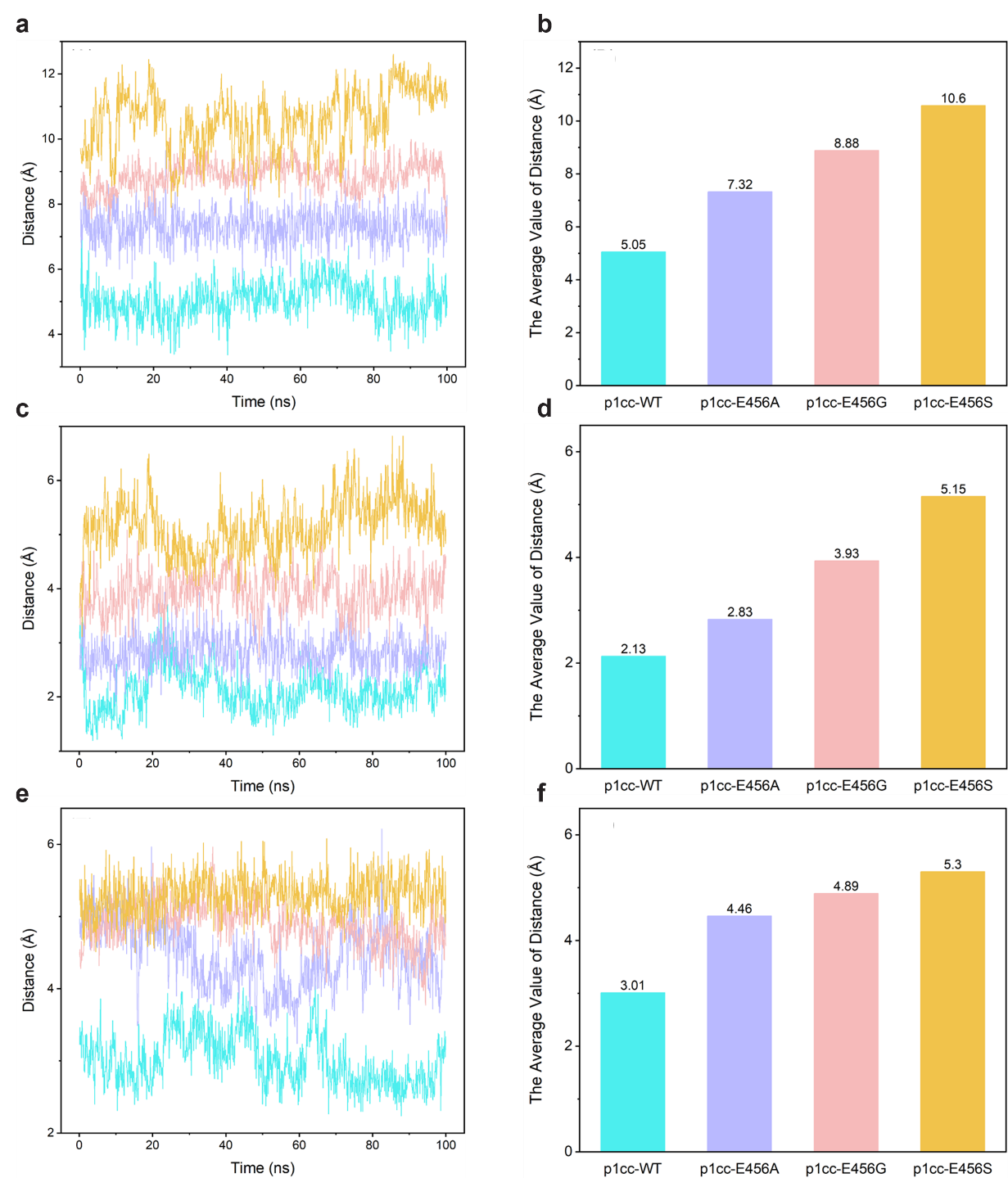

**Figure S8.** MD simulations of WT and variants at the E456 site. **a**, Distance between the substrate’s O–NH₂ group and the WT and three mutant proteins centroid during the MD simulation. **b**, The average distance between the substrate’s O–NH₂ group and the WT and three E456 mutant proteins centroid. **c**, Distance between the substrate centroid and the protein centroid. **d**, The average distance between the substrate centroid and the WT and three E456 mutant proteins centroid. **e**, Distance between the substrate centroid and the centroid of the binding pocket. **f**, The average distance between the substrate centroid and the WT and three E456 mutant proteins centroid. The binding pocket was defined as residues within 4 Å of the initial substrate conformation, including G332, G333, H342, D343, D345, D395, T396, M397, D398, G448, W449, T450, G451, and N473. Line plots in panels **a**, **c**, and **e** represent distance fluctuations during MD simulations, whereas bar charts show the corresponding average values. In panels **b**, **d**, and **f**, blue, purple, pink, and orange denote wild type, E456A, E456G, and E456S, respectively.

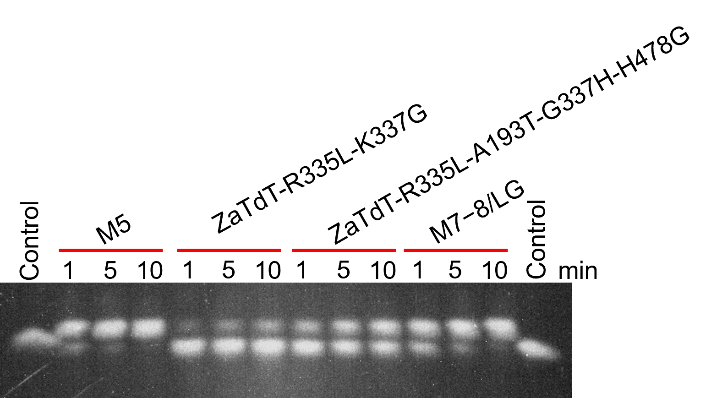

**Figure S9.** Activity comparison of ZaTdT mutants and M5 for P1-CC with 3′-ONH_2_-dATP. Extension conditions: 200 mM 3-(N-morpholino) propanesulfonic acid (MOPS) buffer (pH7.2), 0.05 mg/mL enzyme, 1 µM iDNA P1-CC, 0.2 mM 3′-ONH_2_-dATP, 30 ℃, 1/5/10 min.

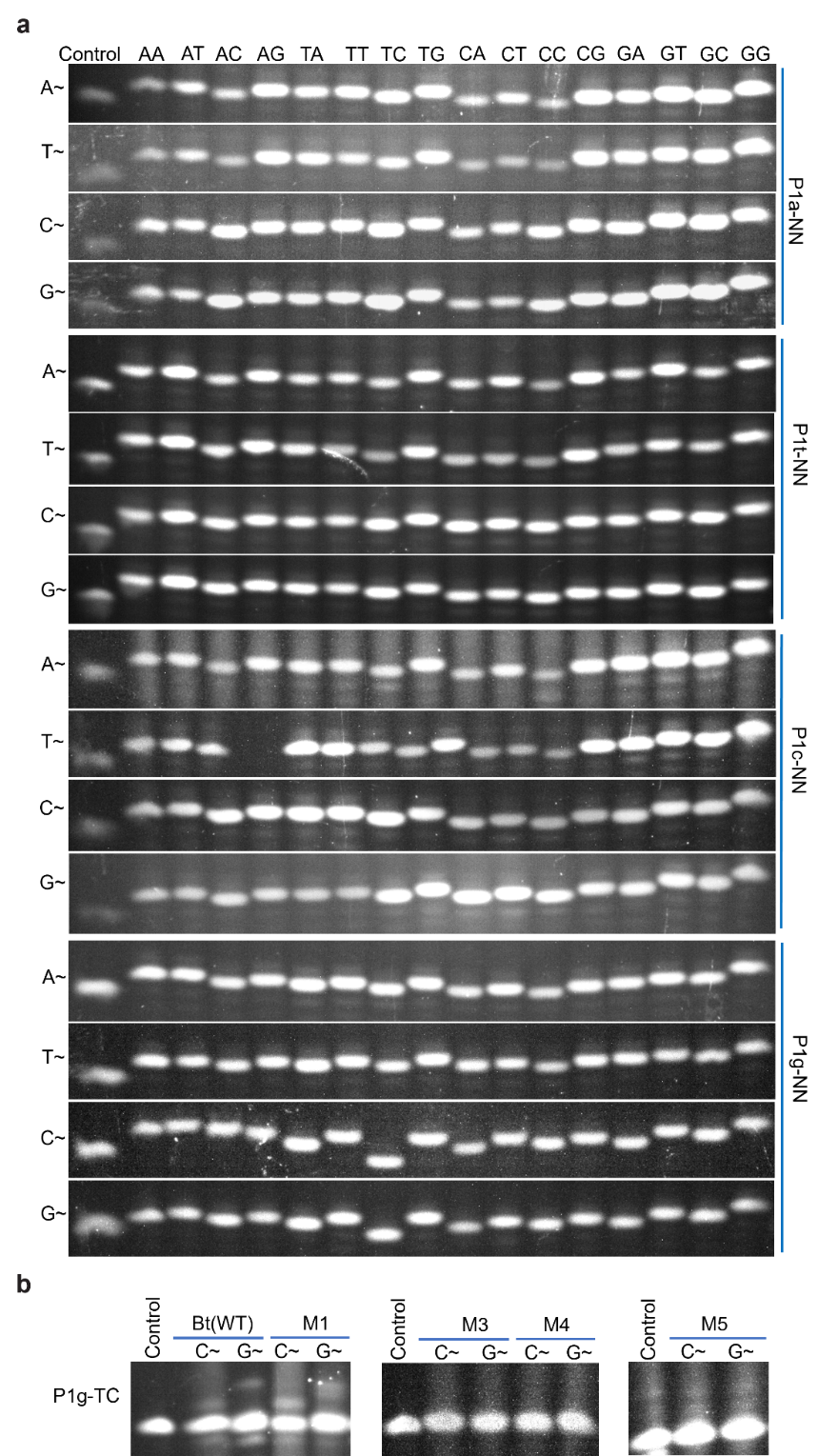

**Figure S10.** Denaturing PAGE analysis. **a**, M5 catalyzing the reactions between 64 (4×4×4) iDNAs containing four trinucleotide combinations at their 3′ end and four 3′-ONH_2_-dNTPs. **b**, BtTdT mutants catalyzing P1g-TC elongating 3′-ONH_2_-dC/GTP. Extension conditions: EDS buffer (15% (v/v) DMSO, 50 mM *O*-benzylhydroxylamine HCl, 10% (v/v) glycerol, 0.05% (v/v) Tween 20, 2.5 mM tris-HCl, 40 mM NaCl, 0.5 mM Hepes, 0.5 M cacodylic acid, and a final pH of 7.0), 1 mg/mL enzyme, 1 µM iDNAs, 0.2 mM 3′-ONH_2_-dATP, 40 ℃, 2 min.

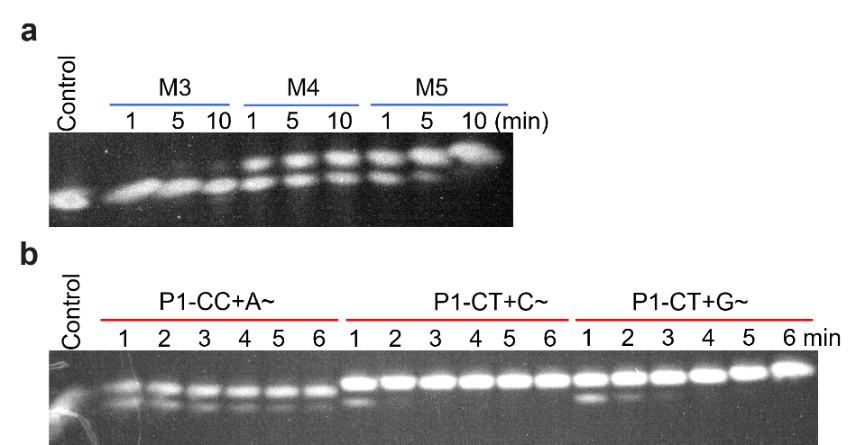

**Figure S11.** Activity comparison of BtTdT mutants for P1-CC with 3′-ONH_2_-dATP. Extension conditions: EDS buffer (pH 7.0), 0.05 mg/mL enzyme, 1 µM iDNA P1-CC, 0.2 mM 3′-ONH_2_-dATP, 30 ℃.

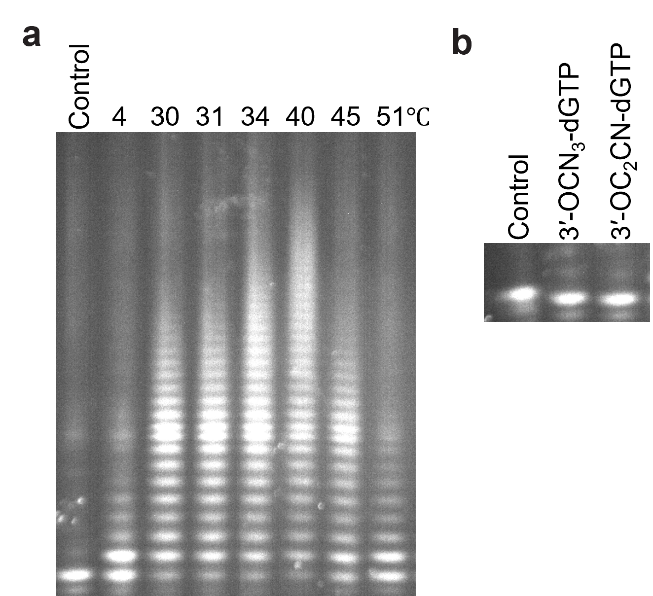

**Figure S12.** Activity of M5. **a**, Optimal reaction temperature of M5. **b**, The incorporation of 3′-OCN_3_ or 3′-OC_2_H_2_CN dGTP for M5. Extension conditions: MOPS buffer (pH 7.0), 0.5 (for **a**) or 1 (for **b**) mg/mL M5, 1 µM iDNA P1-CC, 1 mM dATP (for **a**) or 0.1 mM 3′-OCN_3_ / 3′-OC_2_H_2_CN dGTP (for **b**), 10 min.

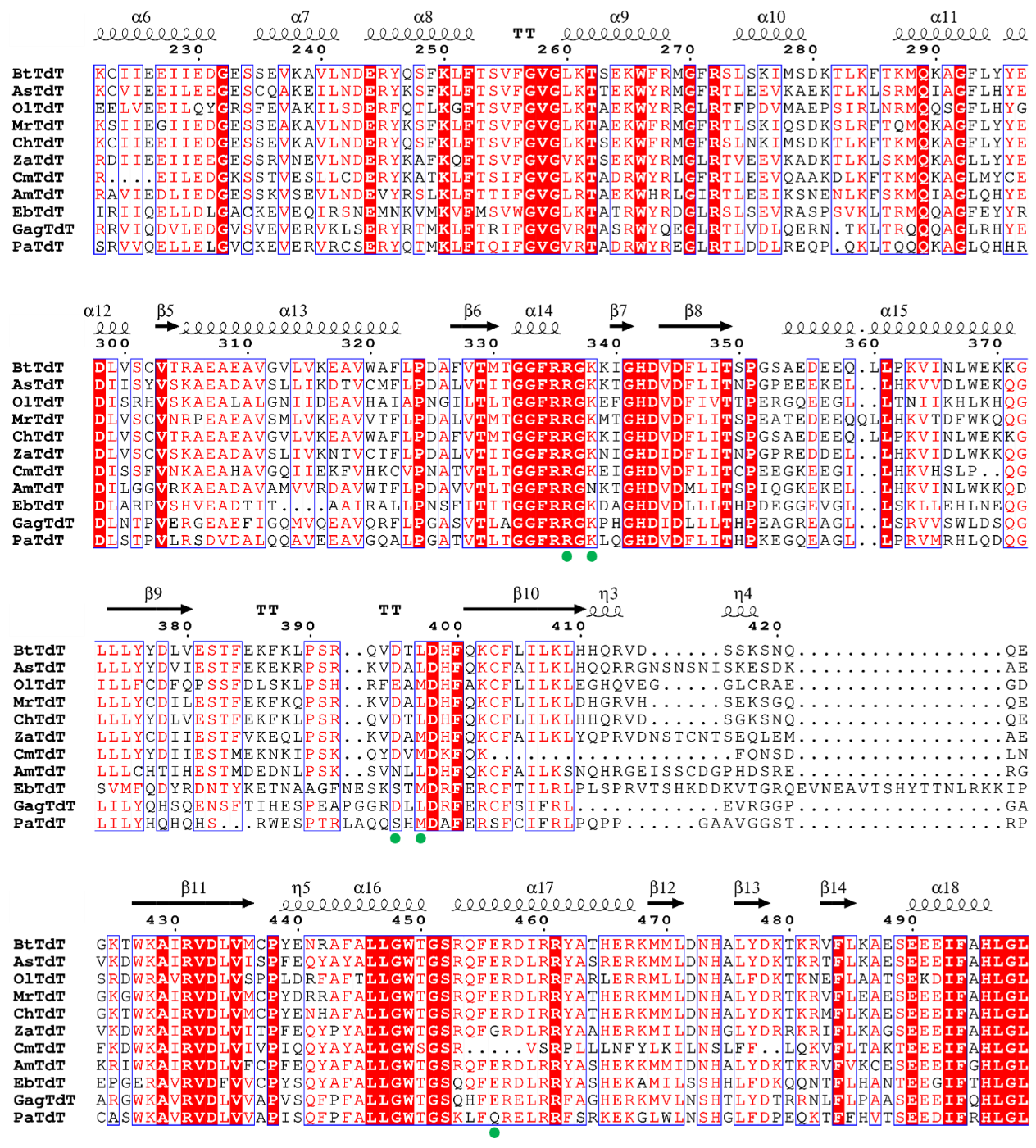

**Figure S13.** Multiple sequence alignment of TdTs, with the five mutated key residues indicated by solid green circles.

**Table S1.** TdT enzymes studied in this work.

| **Protein** | **Species name** | **Class name** | **UniProt or NCBI ID^a^** |
| --- | --- | --- | --- |
| AsTdT | *Alligator sinensis* | Reptilia | A0A3Q0HEY8 |
| BtTdT | *Bos taurus* | Mammalia | P06526 |
| ChTdT | *Capra hircus* | Mammalia | A0A452DR07 |
| CmTdT | *Callorhinchus milii* | Chondrichthyes | A0A4W3JXA8 |
| GagTdT | *Gopherus agassizii* | Reptilia | A0A452HS39 |
| MrTdT | *Mus musculus* | Mammalia | P09838 |
| OlTdT | *Oryzias latipes* | Osteichthyes | A0A3B3IIK7 |
| PaTdT | *Papio anubis* | Mammalia | A0A096NAR8 |
| ZaTdT | *Zonotrichia albicollis* | Aves | XP_026655424.1 (NCBI) |

a: The sequences of the corresponding proteins are given in supplementary table.

**Table S2.** Amino acid sequences of TdTs used in this study.

| **Construct** | **Amino acid sequence** |
| --- | --- |
| AmTdT | MYAFPTTRIAPRRKQPKCIKPPKCTSKYDIKFKDIAIYILERKMGASRRYFLMELARKKGFRVEPDLSEYVTHVVSEKNSGAEVLEWLQAKKAGSIPNVAILDISWFTDCMGAGQPVEIERKHRLTLQKICVCKSPSPVVPSRVGVSQYACQRKTTLDNKNTLFTDAFEILAENYEFRENERSCLSFRQAASVLKSLTFTIAGMADVDGLPGFGDHIRAVIEDLIEDGESSKVSEVLNDEVYRSLKLFTTIFGVGLRTAEKWHRLGIRTLEEIKSNENLKFSKMQIAGLQHYEDILGGVRKAEADAVAMVVRDAVWTFLPDAVVTLTGGFRRGNKTGHDVDMLITSPIQGKEKELLHKVINLWKKQDLLLCHTIHESTMDEDNLPSKSVNLLDHFQKCFAILKSNQHRGEISSCDGPHDSRERGKRIWKAIRVDLVFCPFEQYAFALLGWTGSRQFERDLRRYASHEKKMMIDNHALYDKTKRVFVKCESEEEIFGHLGLEYIDPVERNA |
| AsTdT | MNKAFYQDWEGGKQGDGCQQLGCQVNQQFGKHTRSHEADCAVSMTVQQACSATSETCVSEASSLAADRVSQYACQRRTTLNNYNKKFTDAFEILAENCEFRENRLGCLEFLRAASVLKFLPFPIVKMKNIEGLPCMGDKVKCVIEEILEEGESCQAKEILNDERYKSFKLFTSVFGVGLKTTEKWYRMGFRTLEEVKAEKTLKLSRMQIAGFLHYEDIISYVSKAEADAVSLLIKDTVCMFLPDALVTITGGFRRGKKTGHDVDFLITNPGPEEEKELLHKVVDLWEKQGLLLYYDVIESTFEKEKRPSRKVDALDHFQKCFAILKLHQQRRGNSNSNISKESDKAEVKDWKAIRVDLVISPFEQYAYALLGWTGSRQFERDLRRYASRERKMMLDNHALYDKTKRTFLKAESEEEIFAHLGLDYIEPWERNA |
| BtTdT | MDPLCTASSGPRKKRPRQVGASMASPPHDIKFQNLVLFILEKKMGTTRRNFLMELARRKGFRVENELSDSVTHIVAENNSGSEVLEWLQVQNIRASSQLELLDVSWLIESMGAGKPVEITGKHQLVVRTDYSATPNPGFQKTPPLAVKKISQYACQRKTTLNNYNHIFTDAFEILAENSEFKENEVSYVTFMRAASVLKSLPFTIISMKDTEGIPCLGDKVKCIIEEIIEDGESSEVKAVLNDERYQSFKLFTSVFGVGLKTSEKWFRMGFRSLSKIMSDKTLKFTKMQKAGFLYYEDLVSCVTRAEAEAVGVLVKEAVWAFLPDAFVTMTGGFRRGKKIGHDVDFLITSPGSAEDEEQLLPKVINLWEKKGLLLYYDLVESTFEKFKLPSRQVDTLDHFQKCFLILKLHHQRVDSSKSNQQEGKTWKAIRVDLVMCPYENRAFALLGWTGSRQFERDIRRYATHERKMMLDNHALYDKTKRVFLKAESEEEIFAHLGLDYIEPWERNA |
| ChTdT | MDPLCTASSGPRKKRPRQVGASMASPPHDIKFQNLVLFILEKKMGTTRRNFLMELARRKGFRVENELSDSVTHIVAENNSGSEVLEWLQVQNIRASSQLELLDVSWLIESMGAGKPVEITGKHQLVVRTDYSATPNPGFQKISQYACQRKTTLNNYNHVFTDAFEILAENSEFKENEVSYVTFMRAASVLKSLPFTIISMRDTEGIPCLGDKVKCIIEEIIEDGESSEVKAVLNDERYQSFKLFTSVFGVGLKTSEKWFRMGFRSLNKIMSDKTLKFTKMQKAGFLYYEDLVSCVTRAEAEAVGVLVKEAVWAFLPDAFVTMTGGFRRGKKIGHDVDFLITSPGSAEDEEQLLPKVINLWEKKGLLLYYDLVESTFEKFKLPSRQVDTLDHFQKCFLILKLHHQRVDSGKSNQQEGKTWKAIRVDLVMCPYENHAFALLGWTGSRQFERDIRRYATHERKMMLDNHALYDKTKRMFLKAESEEEIFAHLGLDYIEPWERNA |
| CmTdT | MASTDRLGGTGILPKMKRKKVADSCSHDEYKIKFQGLNIFIVERKMGSTRRTFLMDLARKKGFRVNDKLSDAVTHIVAENNSWNEIWDWLQIHKLSNTNTLEMLDISWFTDSMGAGKPVDIQERHRLFFKKIPLSLRHSWSPVLSCIRPYVQIVFRTGLQQRDALEILAENCEFNENERSYVVFARATSVLKSLPYAISSMTMLEGLPNMEVQPREILEDGKSSTVESLLCDERYKATKLFTSIFGVGLKTADRWYRLGFRTLEEVQAAKDLKFTKMQKAGLMYCEDISSFVNKAEAHAVGQIIEKFVHKCVPNATVTLTGGFRRGKEIGHDVDFLITCPEEGKEEGILHKVHSLPQGLLLYYDIIESTMEKNKIPSKQYDVMDKFQKFQNSDLNFKDWKAIRVDLVIVPIQQYAYALLGWSGSRVSRPLLLNFYLKILNSLFFLQKVFLTAKTEEEIFAHLGLEYIEPWERNA |
| EbTdT | MDPACSTKRRRVGDPGAQRADRPGTKFPEVMIFVRESKMGSSRRAMISALAREKGFGVADSFSTSVTHVVAERNQPKDVWIWLKKVSAKCGLEVLPEVLDISWLTDCMESGCRLPVEDRHRLKDESEEEVCTKSPSIPIYACQRKTPVNHLNHHFTCALEALAEGAEFSESDGRALAFRRAASVLKSLPFRVAKIEQLRGVPCLGKHSIRIIQELLDLGACKEVEQIRSNEMNKVMKVFMSVWGVGLKTATRWYRDGLRSLSEVRASPSVKLTRMQQAGFEYYRDLARPVSHVEADTITAAIRALLPNSFITITGGFRRGKDAGHDVDLLLTHPDEGGEVGLLSKLLEHLNEQGSVMFQDYRDNTYKETNAAGFNESKSTMDRFERCFTILRLPLSPRVTSHKDDKVTGRQEVNEAVTSHYTTNLRKKIPEPGERAVRVDFVVCPYSQYAFALLGWTGSQQFERDLRRYASHEKAMILSSHHLFDKQQNTFLHANTEEGIFTHLGLPYLEPSYRNA |
| GagTdT | MALVPLKRRRRAPSPPAGPLGAGRFPAVVLCLVEKRMGASRRAFLTQLARAKGFRVDGAYSAAVTHVVSEQNSGNEVARWLEQQREECGSGGDPALLDISWFTESMGAGRPVEIESRHRLRVSWAGGSQGMGWGDPTMAPYACQRRTPLLHSNQPLTEVLETLAEEASFSGSEGRSLAFTRAASVLKALPGRLSALEELGPLPGIGEHSRRVIQDVLEDGVSVEVERVKLSERYRTMKLFTRIFGVGVRTASRWYQEGLRTLVDLQERNTKLTRQQQAGLRHYEDLNTPVERGEAEFIGQMVQEAVQRFLPGASVTLAGGFRRGKPHGHDIDLLLTHPEAGREAGLLSRVVSWLDSQGLILYQHSQENSFTIHESPEAPGGRDLLDRFERCFSIFRLEVRGGPGAARGWKAVRVDLVVAPVSQFPFALLGWTGSQHFERELRRFAGHERKMVLNSHTLYDTRRNLFLPAASEEEIFQHLGLEYMPPAQRNA |
| MrTdT | MDPLQAVHLGPRKKRPRQLGTPVASTPYDIRFRDLVLFILEKKMGTTRRAFLMELARRKGFRVENELSDSVTHIVAENNSGSDVLEWLQLQNIKASSELELLDISWLIECMGAGKPVEMMGRHQLVVNRNSSPSPVPGSQNVPAPAVKKISQYACQRRTTLNNYNQLFTDALDILAENDELRENEGSCLAFMRASSVLKSLPFPITSMKDTEGIPCLGDKVKSIIEGIIEDGESSEAKAVLNDERYKSFKLFTSVFGVGLKTAEKWFRMGFRTLSKIQSDKSLRFTQMQKAGFLYYEDLVSCVNRPEAEAVSMLVKEAVVTFLPDALVTMTGGFRRGKMTGHDVDFLITSPEATEDEEQQLLHKVTDFWKQQGLLLYCDILESTFEKFKQPSRKVDALDHFQKCFLILKLDHGRVHSEKSGQQEGKGWKAIRVDLVMCPYDRRAFALLGWTGSRQFERDLRRYATHERKMMLDNHALYDRTKRVFLEAESEEEIFAHLGLDYIEPWERNA |
| OlTdT | MLRWPHPKKRLRPEETTLSRSKKAVFEDVHIFLVERKMGRSRRSFLTQLARSKGFVVEDVLSDAVTHVVSEDSEASSLWAWLKGHSPSDPSKMHVLDISWFTDSMKERRPVAVETKHLIQDVLPEVSTPPSVVAVSQYACQRRTTTENHNKTLTDAFEVLAENYEFNEMEGQCLAFRRAASVLKSLSWQVRSSHQVHDLPCLGETMEELVEEILQYGRSFEVAKILSDERFQTLKGFTSVFGVGLKTAEKWYRRGLRTFPDVMAEPSIRLNRMQQSGFLHYGDISRHVSKAEALALGNIIDEAVHAIAPNGILTLTGGFRRGKEFGHDVDFIVTTPERGQEEGLLTNIIKHLKHQGILLFCDFQPSSFDLSKLPSHRFEAMDHFAKCFLILKLEGHQVEGGLCRAEGDSRDWRAVRVDLVSPPLDRFAFTLLGWTGSRQFERDLRRFARLERRMLLDNHALFDKTKNEFLAATSEKDIFAHLGLEYIEPWQRNA |
| PaTdT | MLPKRRRARVRSPSSDATSSTPPSTRFPGVAIYLVEPRMGRSRRAFLTRLARSKGFRVLDACSSEATHVVMEQTSAEEAVSWQERRMTAAPPGCTPPALLDISWLTESLAAGQPVPVECRHRLEVTGPRKGPLSPAWMPTYACQRPTPLTHHNTSLSEALETLAEAAGFEGSEGRLLTFCRAASVLKALPSPVTTLSQLQGLPHLGEHSSRVVQELLELGVCKEVERVRCSERYQTMKLFTQIFGVGVRTADRWYREGLRTLDDLREQPQKLTQQQKAGLQHHRDLSTPVLRSDVDALQQAVEEAVGQALPGATVTLTGGFRRGKLQGHDVDFLITHPKEGQEAGLLPRVMRHLQDQGLILYHQHQHSRWESPTRLAQQSHMDAFERSFCIFRLPQPPGAAVGGSTRPCASWKAVRVDLVVAPISQFPFALLGWTGSKLFQRELRRFSRKEKGLWLNSHGLFDPEQKTFFHVTSEEDIFRHLGLEYLPPEQRNA |
| ZaTdT | MDRFKAPAVISQRKRQKGLHSPKLSCSYEIKFSNFVIFIMQRKMGLTRRMFLMELGRRKGFRVESELSDSVTHIVAENNSYLEVLDWLKGQAVGDSSRFELLDISWFTACMEAGRPVDSEVKYRLMEQSQSLPLNMPALEMPAFIATKVSQYSCQRKTTLNNYNKKFTDAFEVMAENYEFKENEIFCLEFLRAASLLKSLPFSVTRMKDIQGLPCVGDQVRDIIEEIIEEGESSRVNEVLNDERYKAFKQFTSVFGVGVKTSEKWYRMGLRTVEEVKADKTLKLSKMQKAGLLYYEDLVSCVSKAEADAVSLIVKNTVCTFLPDALVTITGGFRRGKNIGHDIDFLITNPGPREDDELLHKVIDLWKKQGLLLYCDIIESTFVKEQLPSRKVDAMDHFQKCFAILKLYQPRVDNSTCNTSEQLEMAEVKDWKAIRVDLVITPFEQYPYALLGWTGSRQFGRDLRRYAAHERKMILDNHGLYDRRKRIFLKAGSEEEIFAHLGLDYVEPWERNA |
| Bt15AA | MGSSHHHHHHKKISQYACQRKTTLNNYNHIFTDAFEILAENSEFKENEVSYVTFMRAASVLKSLPFTIISMKDTEGIPCLGDKVKCIIEEIIEDGESSEVKAVLNDERYQSFKLFTSVFGVGLKTSEKWFRMGFRSLSKIMSDKTLKFTKMQKAGFLYYEDLVSCVTRAEAEAVGVLVKEAVWAFLPDAFVTMTGGFRRGKKIGHDVDFLITSPGSAEDEEQLLPKVINLWEKKGLLLYYDLVESTFEKFKLPSRQVDTLDHFQKCFLILKLHHQRVDSSKSNQQEGKTWKAIRVDLVMCPYENRAFALLGWTGSRQFERDIRRYATHERKMMLDNHALYDKTKRVFLKAESEEEIFAHLGLDYIEPWERNA |
| M3 | MGSSHHHHHHKKISQYACQRKTTLNNYNHIFTDAFEILAENSEFKENEVSYVTFMRAASVLKSLPFTIISMKDTEGIPCLGDKVKCIIEEIIEDGESSEVKAVLNDERYQSFKLFTSVFGVGLKTSEKWFRMGFRSLSKIMSDKTLKFTKMQKAGFLYYEDLVSCVTRAEAEAVGVLVKEAVWAFLPDAFVTMTGGFRLGGKIGHDVDFLITSPGSAEDEEQLLPKVINLWEKKGLLLYYDLVESTFEKFKLPSRQVDTMDHFQKCFLILKLHHQRVDSSKSNQQEGKTWKAIRVDLVMCPYENRAFALLGWTGSRQFERDIRRYATHERKMMLDNHALYDKTKRVFLKAESEEEIFAHLGLDYIEPWERNA |
| M4 | MGSSHHHHHHKKISQYACQRKTTLNNYNHIFTDAFEILAENSEFKENEVSYVTFMRAASVLKSLPFTIISMKDTEGIPCLGDKVKCIIEEIIEDGESSEVKAVLNDERYQSFKLFTSVFGVGLKTSEKWFRMGFRSLSKIMSDKTLKFTKMQKAGFLYYEDLVSCVTRAEAEAVGVLVKEAVWAFLPDAFVTMTGGFRLGGKIGHDVDFLITSPGSAEDEEQLLPKVINLWEKKGLLLYYDLVESTFEKFKLPSRQVDTMDHFQKCFLILKLHHQRVDSSKSNQQEGKTWKAIRVDLVMCPYENRAFALLGWTGSRQFSRDIRRYATHERKMMLDNHALYDKTKRVFLKAESEEEIFAHLGLDYIEPWERNA |
| M5 | MGSSHHHHHHKKISQYACQRKTTLNNYNHIFTDAFEILAENSEFKENEVSYVTFMRAASVLKSLPFTIISMKDTEGIPCLGDKVKCIIEEIIEDGESSEVKAVLNDERYQSFKLFTSVFGVGLKTSEKWFRMGFRSLSKIMSDKTLKFTKMQKAGFLYYEDLVSCVTRAEAEAVGVLVKEAVWAFLPDAFVTMTGGFRLGGKIGHDVDFLITSPGSAEDEEQLLPKVINLWEKKGLLLYYDLVESTFEKFKLPSRQVGTMDHFQKCFLILKLHHQRVDSSKSNQQEGKTWKAIRVDLVMCPYENRAFALLGWTGSRQFSRDIRRYATHERKMMLDNHALYDKTKRVFLKAESEEEIFAHLGLDYIEPWERNA |
| M7-8/LG | MGSSHHHHHHMKVSQYACQRRTTLNNHNKRFTDAFEIMAEYYEFNENEGRCLAFRRAASVLKSLPFTVTRMKDIQGLPCFGDHVRRIIQEILEHGESSEVERVLNDERYQAFKLFTSVFGVGVKTAEKWYRMGLRTVEEVKADKTLKLTKMQKAGLQYYEDLVSCVSKAEADAISQIVKETVWAFLPDALVTMTGGFRLGGEIGHDVDFLITNPGPREDDELLHKVIDLWKKQGLLLYCDIIESTFDKSKLPSRKVDAMDHFQKCFCILKLYQPRVDNSTYNTSKQLDMAEVKDWKAVRVDLVVTPYEQYAFALLGWTGSKQFNRDLRRYARHERKMLLDNHGLYDRTQKIFLKATSEEEIFAHLGLEYIPPWERNA |
| ZaTdT-R335L-A193T-G337H-H478G | MGSSHHHHHHMDRFKAPAVISQRKRQKGLHSPKLSCSYEIKFSNFVIFIMQRKMGLTRRMFLMELGRRKGFRVESELSDSVTHIVAENNSYLEVLDWLKGQAVGDSSRFELLDISWFTACMEAGRPVDSEVKYRLMEQSQSLPLNMPALEMPAFIATKVSQYSCQRKTTLNNYNKKFTDAFEVMAENYEFKENEIFCLEFLRTASLLKSLPFSVTRMKDIQGLPCVGDQVRDIIEEIIEEGESSRVNEVLNDERYKAFKQFTSVFGVGVKTSEKWYRMGLRTVEEVKADKTLKLSKMQKAGLLYYEDLVSCVSKAEADAVSLIVKNTVCTFLPDALVTITGGFRLGHNIGHDIDFLITNPGPREDDELLHKVIDLWKKQGLLLYCDIIESTFVKEQLPSRKVDAMDHFQKCFAILKLYQPRVDNSTCNTSEQLEMAEVKDWKAIRVDLVITPFEQYPYALLGWTGSRQFGRDLRRYAAHERKMILDNGGLYDRRKRIFLKAGSEEEIFAHLGLDYVEPWERNA |
| ZaTdT-R335L-K337G | MGSSHHHHHHMDRFKAPAVISQRKRQKGLHSPKLSCSYEIKFSNFVIFIMQRKMGLTRRMFLMELGRRKGFRVESELSDSVTHIVAENNSYLEVLDWLKGQAVGDSSRFELLDISWFTACMEAGRPVDSEVKYRLMEQSQSLPLNMPALEMPAFIATKVSQYSCQRKTTLNNYNKKFTDAFEVMAENYEFKENEIFCLEFLRAASLLKSLPFSVTRMKDIQGLPCVGDQVRDIIEEIIEEGESSRVNEVLNDERYKAFKQFTSVFGVGVKTSEKWYRMGLRTVEEVKADKTLKLSKMQKAGLLYYEDLVSCVSKAEADAVSLIVKNTVCTFLPDALVTITGGFRLGGNIGHDIDFLITNPGPREDDELLHKVIDLWKKQGLLLYCDIIESTFVKEQLPSRKVDAMDHFQKCFAILKLYQPRVDNSTCNTSEQLEMAEVKDWKAIRVDLVITPFEQYPYALLGWTGSRQFGRDLRRYAAHERKMILDNHGLYDRRKRIFLKAGSEEEIFAHLGLDYVEPWERNA |

**Table S3.** Vectors used in this study.

| **Vectors** | **Description** | **Source** |
| --- | --- | --- |
| pETDuet-His_6_-AsTdT | Expression vector of N-terminal His_6_-tag AsTdT | this study |
| pETDuet-His_6_-BtTdT | Expression vector of N-terminal His_6_-tag BtTdT | this study |
| pETDuet-His_6_-ChTdT | Expression vector of N-terminal His_6_-tag ChTdT | this study |
| pETDuet-His_6_-CmTdT | Expression vector of N-terminal His_6_-tag CmTdT | this study |
| pETDuet-His_6_-GagTdT | Expression vector of N-terminal His_6_-tag GagTdT | this study |
| pETDuet-His_6_-MrTdT | Expression vector of N-terminal His_6_-tag MrTdT | this study |
| pETDuet-His_6_-OlTdT | Expression vector of N-terminal His_6_-tag OlTdT | this study |
| pETDuet-His_6_-PaTdT | Expression vector of N-terminal His_6_-tag PaTdT | this study |
| pETDuet-His_6_-ZaTdT | Expression vector of N-terminal His_6_-tag ZaTdT | this study |
| pETDuet-His_6_-ZaTdT-R335L-A193T-G337H-H478G | Expression vector of N-terminal His_6_-tag ZaTdT-R335L-A193T-G337H-H478G | this study |
| pETDuet-His_6_-ZaTdT-R335L-K337G | Expression vector of N-terminal His_6_-tag ZaTdT-R335L-K337G | this study |
| pETDuet-His_6_-M7-8/LG | Expression vector of N-terminal His_6_-tag M7-8/LG | this study |

**Table S4.** Primers used in this study.

| **Primers** | **Sequence 5´ to 3´** | **Description** |
| --- | --- | --- |
| Bt0AA-F | CAGCAGCCATCACCATCATCACCACAACTATAACCATATTTTCACCG | For cloning Bt0AA |
| Bt2AA-F | CAGCAGCCATCACCATCATCACCACCTGAACAACTATAACCATATTT | For cloning Bt2AA |
| Bt5AA-F | CAGCAGCCATCACCATCATCACCACAAAACCACCCTGAACAACTATA | For cloning Bt5AA |
| Bt10AA-F | CAGCAGCCATCACCATCATCACCACTATGCGTGCCAGCGCAAAACCA | For cloning Bt10AA |
| Bt15AA-F | CAGCAGCCATCACCATCATCACCACAAAAAGATTAGCCAGTATGCGT | For cloning Bt15AA |
| Bt20AA-F | CAGCAGCCATCACCATCATCACCACCCGCCGCTGGCGGTTAAAAAGA | For cloning Bt20AA |
| Bt25AA-F | CAGCAGCCATCACCATCATCACCACGGTTTTCAGAAAACCCCGCCGC | For cloning Bt25AA |
| Bt30AA-F | CAGCAGCCATCACCATCATCACCACGCGACCCCGAATCCGGGTTTTC | For cloning Bt30AA |
| Bt37AA-F | CAGCAGCCATCACCATCATCACCACGTGGTGCGCACCGATTATAGCG | For cloning Bt37AA |
| Bt-R | CTGTTCGACTTAAGCATTATGCTTACGCGTTGCGTTCCCACGGTTC | For cloning Bt wild type or mutants |
| P0 | TTTTTTTTTTTTTTTTTTTT | Substrate iDNA |
| P1 | TAATACGACTCACTA | Substrate iDNA |
| P2 | biotin-ACTAGGACGACTCGAATT | Substrate iDNA |
| P1-AA | TAATACGACTCACTAAA | Substrate iDNA |
| P1-AT | TAATACGACTCACTAAT | Substrate iDNA |
| P1-AC | TAATACGACTCACTAAC | Substrate iDNA |
| P1-AG | TAATACGACTCACTAAG | Substrate iDNA |
| P1-TA | TAATACGACTCACTATA | Substrate iDNA |
| P1-TT | TAATACGACTCACTATT | Substrate iDNA |
| P1-TC | TAATACGACTCACTATC | Substrate iDNA |
| P1-TG | TAATACGACTCACTATG | Substrate iDNA |
| P1-CA | TAATACGACTCACTACA | Substrate iDNA |
| P1-CT | TAATACGACTCACTACT | Substrate iDNA |
| P1-CC | TAATACGACTCACTACC | Substrate iDNA |
| P1-CG | TAATACGACTCACTACG | Substrate iDNA |
| P1-GA | TAATACGACTCACTAGA | Substrate iDNA |
| P1-GT | TAATACGACTCACTAGT | Substrate iDNA |
| P1-GC | TAATACGACTCACTAGC | Substrate iDNA |
| P1-GG | TAATACGACTCACTAGG | Substrate iDNA |
| P1t-AA | TAATACGACTCACtAA | Substrate iDNA |
| P1t-AT | TAATACGACTCACtAT | Substrate iDNA |
| P1t-AC | TAATACGACTCACtAC | Substrate iDNA |
| P1t-AG | TAATACGACTCACtAG | Substrate iDNA |
| P1t-TA | TAATACGACTCACtTA | Substrate iDNA |
| P1t-TT | TAATACGACTCACtTT | Substrate iDNA |
| P1t-TC | TAATACGACTCACtTC | Substrate iDNA |
| P1t-TG | TAATACGACTCACtTG | Substrate iDNA |
| P1t-CA | TAATACGACTCACtCA | Substrate iDNA |
| P1t-CT | TAATACGACTCACtCT | Substrate iDNA |
| P1t-CC | TAATACGACTCACtCC | Substrate iDNA |
| P1t-CG | TAATACGACTCACtCG | Substrate iDNA |
| P1t-GA | TAATACGACTCACtGA | Substrate iDNA |
| P1t-GT | TAATACGACTCACtGT | Substrate iDNA |
| P1t-GC | TAATACGACTCACtGC | Substrate iDNA |
| P1t-GG | TAATACGACTCACtGG | Substrate iDNA |
| P1a-AA | TAATACGACTCACaAA | Substrate iDNA |
| P1a-AT | TAATACGACTCACaAT | Substrate iDNA |
| P1a-AC | TAATACGACTCACaAC | Substrate iDNA |
| P1a-AG | TAATACGACTCACaAG | Substrate iDNA |
| P1a-TA | TAATACGACTCACaTA | Substrate iDNA |
| P1a-TT | TAATACGACTCACaTT | Substrate iDNA |
| P1a-TC | TAATACGACTCACaTC | Substrate iDNA |
| P1a-TG | TAATACGACTCACaTG | Substrate iDNA |
| P1a-CA | TAATACGACTCACaCA | Substrate iDNA |
| P1a-CT | TAATACGACTCACaCT | Substrate iDNA |
| P1a-CC | TAATACGACTCACaCC | Substrate iDNA |
| P1a-CG | TAATACGACTCACaCG | Substrate iDNA |
| P1a-GA | TAATACGACTCACaGA | Substrate iDNA |
| P1a-GT | TAATACGACTCACaGT | Substrate iDNA |
| P1a-GC | TAATACGACTCACaGC | Substrate iDNA |
| P1a-GG | TAATACGACTCACaGG | Substrate iDNA |
| P1c-AA | TAATACGACTCACcAA | Substrate iDNA |
| P1c-AT | TAATACGACTCACcAT | Substrate iDNA |
| P1c-AC | TAATACGACTCACcAC | Substrate iDNA |
| P1c-AG | TAATACGACTCACcAG | Substrate iDNA |
| P1c-TA | TAATACGACTCACcTA | Substrate iDNA |
| P1c-TT | TAATACGACTCACcTT | Substrate iDNA |
| P1c-TC | TAATACGACTCACcTC | Substrate iDNA |
| P1c-TG | TAATACGACTCACcTG | Substrate iDNA |
| P1c-CA | TAATACGACTCACcCA | Substrate iDNA |
| P1c-CT | TAATACGACTCACcCT | Substrate iDNA |
| P1c-CC | TAATACGACTCACcCC | Substrate iDNA |
| P1c-CG | TAATACGACTCACcCG | Substrate iDNA |
| P1c-GA | TAATACGACTCACcGA | Substrate iDNA |
| P1c-GT | TAATACGACTCACcGT | Substrate iDNA |
| P1c-GC | TAATACGACTCACcGC | Substrate iDNA |
| P1c-GG | TAATACGACTCACcGG | Substrate iDNA |
| P1g-AA | TAATACGACTCACgAA | Substrate iDNA |
| P1g-AT | TAATACGACTCACgAT | Substrate iDNA |
| P1g-AC | TAATACGACTCACgAC | Substrate iDNA |
| P1g-AG | TAATACGACTCACgAG | Substrate iDNA |
| P1g-TA | TAATACGACTCACgTA | Substrate iDNA |
| P1g-TT | TAATACGACTCACgTT | Substrate iDNA |
| P1g-TC | TAATACGACTCACgTC | Substrate iDNA |
| P1g-TG | TAATACGACTCACgTG | Substrate iDNA |
| P1g-CA | TAATACGACTCACgCA | Substrate iDNA |
| P1g-CT | TAATACGACTCACgCT | Substrate iDNA |
| P1g-CC | TAATACGACTCACgCC | Substrate iDNA |
| P1g-CG | TAATACGACTCACgCG | Substrate iDNA |
| P1g-GA | TAATACGACTCACgGA | Substrate iDNA |
| P1g-GT | TAATACGACTCACgGT | Substrate iDNA |
| P1g-GC | TAATACGACTCACgGC | Substrate iDNA |
| P1g-GG | TAATACGACTCACgGG | Substrate iDNA |

(1)	Salomon‐Ferrer, R.; Case, D. A.; Walker, R. C. An Overview of the Amber Biomolecular Simulation Package. *WIREs Comput. Mol. Sci.* **2013**, *3* (2), 198–210.

(2)	Maier, J. A.; Martinez, C.; Kasavajhala, K.; Wickstrom, L.; Hauser, K. E.; Simmerling, C. ff14SB: Improving the Accuracy of Protein Side Chain and Backbone Parameters from ff99SB. *J. Chem. Theory Comput.* **2015**, *11* (8), 3696–3713.

(3)	Galindo-Murillo, R.; Robertson, J. C.; Zgarbová, M.; Šponer, J.; Otyepka, M.; Jurečka, P.; Cheatham, T. E. Assessing the Current State of Amber Force Field Modifications for DNA. *J. Chem. Theory Comput.* **2016**, *12* (8), 4114–4127.

(4)	Wang, J.; Wolf, R. M.; Caldwell, J. W.; Kollman, P. A.; Case, D. A. Development and Testing of a General Amber Force Field. *J. Comput. Chem.* **2004**, *25* (9), 1157–1174.

(5)	Doshi, U.; Hamelberg, D. Extracting Realistic Kinetics of Rare Activated Processes from Accelerated Molecular Dynamics Using Kramers’ Theory. *J. Chem. Theory Comput.* **2011**, *7* (3), 575–581.

(6)	Lin, Y.; Pan, D.; Li, J.; Zhang, L.; Shao, X. Application of Berendsen Barostat in Dissipative Particle Dynamics for Nonequilibrium Dynamic Simulation. *J. Chem. Phys.* **2017**, *146* (12), 124108.

(7)	Elber, R.; Ruymgaart, A. P.; Hess, B. SHAKE Parallelization. *Eur. Phys. J. Spec. Top.* **2011**, *200* (1), 211–223.

(8)	Simmonett, A. C.; Brooks, B. R. A Compression Strategy for Particle Mesh Ewald Theory. *J. Chem. Phys.* **2021**, *154* (5), 054112.

(9)	Humphrey, W.; Dalke, A.; Schulten, K. VMD: Visual Molecular Dynamics. *J. Mol. Graph.* **1996**, *14* (1), 33–38.

(10) DeLano, W. L. *The PyMOL Molecular Graphics System*, Version 2.3; Schrödinger LLC, 2020.
